# Does rehearsal matter? Left anterior temporal alpha and theta band changes correlate with the beneficial effects of rehearsal on working memory

**DOI:** 10.1101/753350

**Authors:** Chelsea Reichert Plaska, Kenneth Ng, Timothy M. Ellmore

## Abstract

Rehearsal during working memory (WM) maintenance facilitates retrieval. Less is known about how rehearsal modulates WM delay activity. In the present study, 44 participants completed a Sternberg Task with either intact novel scenes or phase-scrambled scenes, which had similar color and spatial frequency but lacked semantic content. During each condition participants generated a descriptive label and covertly rehearsed or suppressed (repeated “the”) during the delay. This was easy in the former but more difficult in the later condition where scenes lacked semantic content. Behavioral performance and EEG delay activity was analyzed as a function of maintenance strategy. Performance during WM revealed a benefit of rehearsal for phase-scrambled but not intact scenes. Examination of the absolute amplitude revealed three underlying sources of activity for rehearsal, including the left anterior temporal (TAL), left and midline parietal regions. Increases in alpha and theta activity in TAL were correlated with improvement in performance on WM with rehearsal only when labeling was not automatic (i.e. phase-scrambled scenes), which may reflect differences in labeling and rehearsal (i.e. semantic associations vs. shallow labels). We conclude that rehearsal only benefits memory for visual stimuli that lack semantic information, and that this is correlated with changes in alpha and theta rhythms.

## Introduction

It is well documented that maintenance via rehearsal benefits memory for verbal stimuli, such as words and numbers, typically via cumulative rehearsal when confronted with a list of verbal stimuli to remember. Baddeley ^1^ established the idea that rehearsal benefits memory, suggesting that the repetition of the to-be-remembered item will refresh the memory trace via the articulatory process in the phonological loop. Studies support that blocking rehearsal with articulatory suppression, repeating a word such as “the” over and over again, decreases performance compared with when someone rehearses ^2,3^. Visual stimuli may also benefit from rehearsal when the stimulus is recoded into a verbal representation ^4-6^. However, the benefit of labeling and rehearsal may depend on the working memory task ^7-9^.

Most visual stimuli, no matter how simple or complex, have an affiliated word representation (i.e. a picture of an ocean with sand and trees has a label of “beach”) which automatically links the stimulus with stored semantic representations ^10^ that contains many features (i.e. sand and ocean) and associated knowledge (i.e. beaches are located in warm places and have palm trees). Thus, combining visual information with a semantic verbal representation may result in a deeper level of encoding ^11,12^ because the stimulus is encoded in both the visual and verbal domains ^13^. The novel features of the visual stimuli may also help to guide attention to salient features which facilitates encoding ^13,14^. Few studies have examined whether complex visual stimuli benefit from recoding, including abstract shapes ^15,16^ and patterns ^4^. Complex stimuli may not benefit from recoding or rehearsal, especially if features lack concreteness such as nameable colors or line orientation ^6^.

In order to understand the mechanisms that support maintenance of visual information, it is critical to examine neural activity during the delay period ^17^ as it supports successful retrieval from working memory. Delay activity has been defined as a period of increased and sustained activation throughout the delay period ^17,18^ and has been identified in prefrontal and posterior parietal regions. Similar to the behavioral rehearsal literature, the delay activity literature has identified these patterns of maintenance with verbal and simple visual stimuli, with few studies measuring delay activity during maintenance of complex visual stimuli. Additionally, few studies of delay activity have controlled the maintenance strategy used by participants during the maintenance of information which is surprising because maintenance strategies can differentially engage brain regions ^19^.

This traditional view of delay activity suggests that it is supported by persistent neuronal firing, which maintains information online until a response is made ^20^. Different types of maintenance strategies may be used by the participant especially without explicit instructions. Rehearsal engages language areas ^21^, especially for verbal stimuli ^22^. More specifically, studies that control for maintenance have identified left temporal region activation reflects the active use of the phonological loop during rehearsal ^2,22^. Activation in the parietal region has been suggested as the store for visual information during maintenance and may serve as the buffer in which information lives until it is needed for retrieval, analogous to the verbal information store ^22^. In fact, it has been proposed that the visual cache, which holds non-spatial visual information, may overlap with the functions of the episodic buffer proposed in the multicomponent model ^10^. The episodic buffer is believed to serve as the interface between the information in the visual and verbal stores with stored semantic knowledge ^23^. Thus, the lateral posterior parietal cortex could represent the area in the brain in which the recoded verbal label is associated with the visually stored picture and the long-term semantic associations, consistent with the output buffer hypothesis ^23,24^. Although, recoding has been proposed to automatically occur with verbalizable stimuli ^25^ and rehearsal ^26^, other mechanisms may support maintenance such as attentional mechanisms that engage a different set of brain regions. Attentional refreshing, which involves directing attention inward to selectively keep information active and prevent interference from other brain regions, engages attentional mechanisms ^27^.

If maintenance strategies are not explicitly controlled, delay period activity is difficult to interpret and may explain recent challenges to the established patterns ^17,28^. Controlling for these components is critical to identifying brain regions that support memory for visual information and how activity in these regions changes when words and semantic meaning are associated with visual information to support memory. In the present study two experiments were completed to understand how articulatory rehearsal influences short- and long-term memory for complex visual stimuli and the underlying delay activity. It was hypothesized that controlling for the maintenance strategy by comparing rehearsal to articulatory suppression would result in differences in behavioral performance and delay period activity. Regardless of the type of complex visual stimulus, it was predicted that rehearsal would provide a behavioral advantage over suppression both in tests of working memory and immediate long-term memory. Delay activity during rehearsal would be sustained throughout the delay period and correlated with performance.

## Results

### Experiment 1

#### Behavioral

Performance on the WM task revealed that there was no significant difference between rehearsal and suppression (.95 vs. .95 proportion correct, *t*_(28)_ = .70, *p* = .49, *d* = .13), suggesting that rehearsal did not provide a short-term behavioral advantage (**Figure 2a and b**). Similarly, there was no long-term behavioral advantage on the delayed recognition task for rehearsal vs. suppression (.80 vs. .78 proportion correct, *t*_(28)_ = 1.38, *p* = .18, *d* = .23).

**Figure 1.**
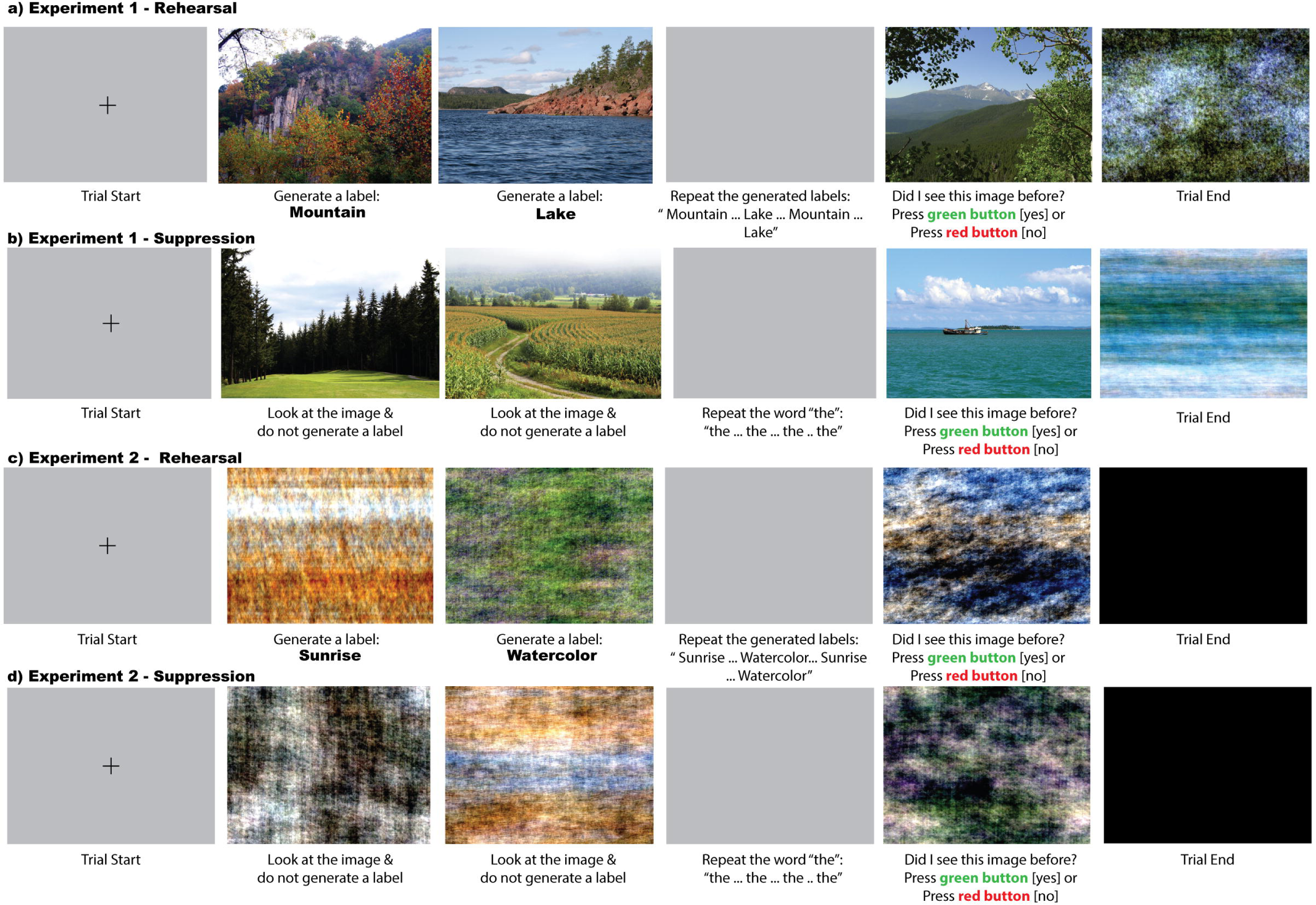
Example Working Memory Trial in Experiment 1 (Intact Scenes) and Experiment 2 (Scrambled Scenes). The task consisted of a low cognitive load (2 images). Participants were presented with a fixation cross (1 sec) that indicates the start of the trial, followed by 2 images in succession (2 sec each), a blank screen during the delay period (6 sec), a probe choice (2 sec), which is either one of the earlier presented images or a new image, and a phase-scrambled image (1 sec) that indicates the end of the trial. An example of a rehearsal trial in which participants generate the label for each image and rehearse during the delay period (a) and an example of a suppression trial in which participants suppress during the delay period (b). Experiment 2 used the same task used in Experiment 1 except with phase-scrambled scenes. An example of a rehearsal trial in which participants generate the label for each phase-scrambled image and rehearse during the delay period (c) and an example of a suppression trial in which participants suppress during the delay period (d).

**Figure 2.**
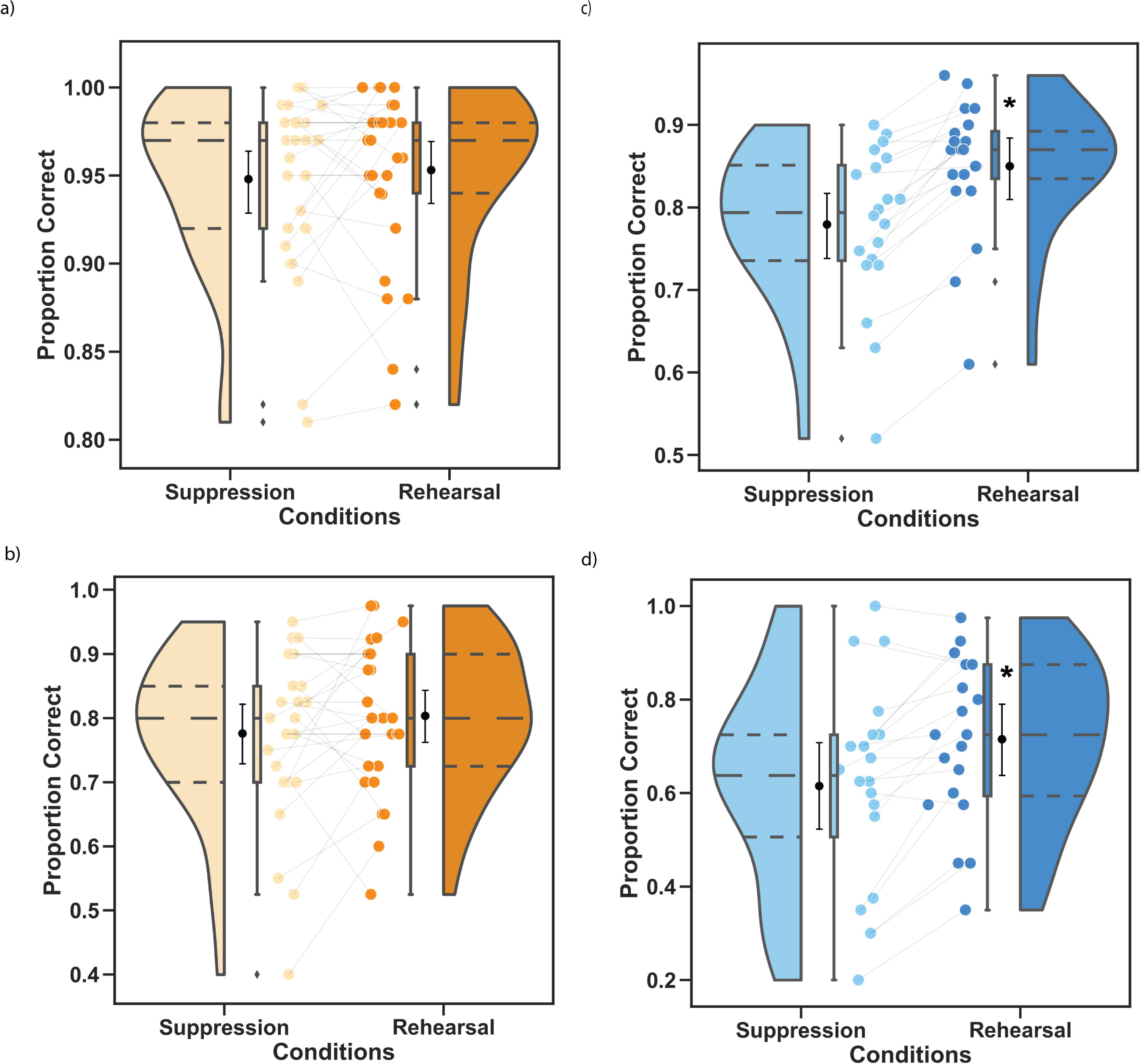
Working Memory and Recognition Performance in Experiment 1 (Intact Scenes) and Experiment 2 (Phase-Scrambled Scenes) Suggests Rehearsal Only Benefits Memory for Scenes that Lack Semantic Content. Performance is shown with four plots for each condition and experiment. 1) The violin plot shows the distribution of performance across each condition; the cluster of similar scores is reflected at the wider (top) portion of the violin and the thinner (bottom) portion with fewer scores. The dashed lines reflect the quartiles of the overall distribution similar to the box plot. 2) The boxplot also shows the distribution of performance for each condition (i.e. the solid lines inside the box align with the dashed lines of the violin plot, representing the quartiles). In addition, the boxplot contains whiskers extend to the minimum and maximum score (1.5 times the median) for each condition and the diamonds reflect outliers (greater than 1.5 times the median). Outliers were included in all analyses. 3) In between the violin plot and boxplot is a point plot, which reflects the mean of performance accuracy, with error bars reflecting the 95% confidence intervals; the confidence intervals were generated with bootstrapping using 1,000 iterations. 4) Finally, the individual dots on the inner-most portion represents single participants score for a condition. The gray line connects the dots that represents performance for the corresponding condition (i.e. performance on suppression (left) with performance on rehearsal (right)). a) Comparison of performance on WM task for Experiment 1 (*n* = 29)., as measured by proportion correct, shows that rehearsal (light orange) provided no benefit for short-term memory as compared with suppression (dark orange on the working memory task (*p* = .49). b) Comparison of performance on the recognition task for Experiment 1, for images from the rehearsal condition (light orange) versus images from the suppression condition (dark orange), suggested that there was no long-term benefit of rehearsal (*p* = .18). c) Examination of performance on the working memory task for Experiment 2 (*n* = 20) shows that rehearsal (light blue) provided a short-term advantage as compared with suppression (dark blue) on the working memory task (*p* < .001). d) Comparison of performance on the recognition task for Experiment 2 also revealed a long-term advantage for images from the rehearsal condition (light blue) versus images from the suppression condition (dark blue), (*p* < .001). These cluster details are described in *Supplementary Table 1*.

#### EEG

Change in sensor-level amplitude over time (temporal spectral analysis) reveals a transient pattern of delay activity during rehearsal (**Figure 3a**). Activity was increased and synchronous early in the delay period in the upper alpha and lower beta range (500 msec to 3000 msec) and became desynchronous later in the delay period (4000 msec to 6000 msec). Temporal spectral analysis revealed no significant difference between rehearsal versus suppression (*p* = .08), nor was rehearsal significantly correlated with working memory (*p* = .46) or recognition performance (*p* = .28). This would suggest that the early pattern of event-related synchronization or enhancement (ERS) in alpha and beta and the late pattern of event-related synchronization or suppression (ERD) in the same frequency bands, was the same regardless of the maintenance mechanism.

**Figure 3.**
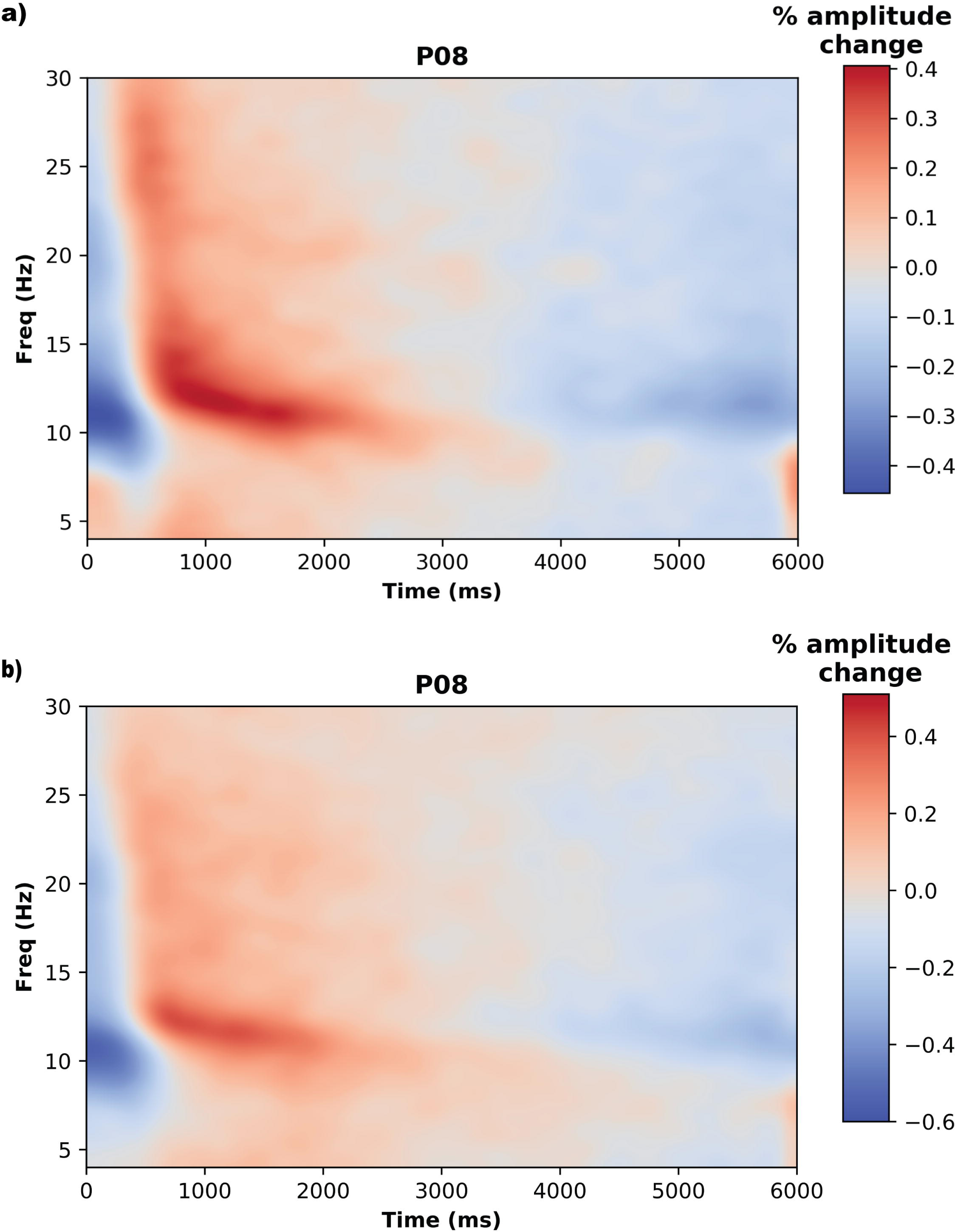
Delay Period Time Frequency Analysis during Rehearsal for Experiment 1 (Intact Scenes) and Experiment 2 (Phase-Scrambled Scenes) Reveal a Transient Pattern of Delay Activity. Time Frequency Analysis plot for the PO8 electrode from the rehearsal condition during the 6-sec delay period in a whole window analysis. Electrode selected based on results from previous research in our lab highlighting the right parieto-occipital region ^52^. a) Time Frequency Analysis for the PO8 electrode in the Rehearsal condition in Experiment 1. b) Time Frequency Analysis for the PO8 electrode in the Rehearsal condition in Experiment 1. Review of the overall pattern of delay activity during rehearsal reveals a similar transient pattern of delay activity for the parieto-occipital censors in Experiments 1 and 2 with an early period of increased synchronous activity in the upper alpha and beta (500 msec to 3000 msec) followed by a period of desynchronous activity in the same frequency bands (4000 msec to 6000 msec). An exploratory analysis of TFA between experiments 1 and 2 (see *Supplementary Results*) revealed that there were no significantly different clusters of brain activity between the conditions. This suggests that regardless of stimulus type (intact scenes or phase-scrambled scenes) a transient pattern of delay activity supports visual working memory.

Sensor-level changes in absolute amplitudes between the two conditions (n=24 subjects) with corrections for multiple comparisons revealed 100 significant clusters (**Supplementary Table 1**, p < .05). Sensor-level analysis revealed two distinct patterns of activity for the rehearsal and suppression conditions (**Supplementary Figure 1**). For the rehearsal condition (**Figure 4b** – P8 electrode - orange clusters), amplitude was greater in the theta and beta range for the left frontal, bilateral fronto-temporal, and central regions, and in the beta range for the right parietal region, throughout the delay period. For the suppression condition (**Figure 4c** – F1 electrode - blue clusters), the amplitude was greater in the upper alpha and lower beta range in the mid-frontal regions early in the delay, and in the theta and upper alpha range in the midline and centro-frontal, right parietal, and occipital regions later in the delay.

**Figure 4.**
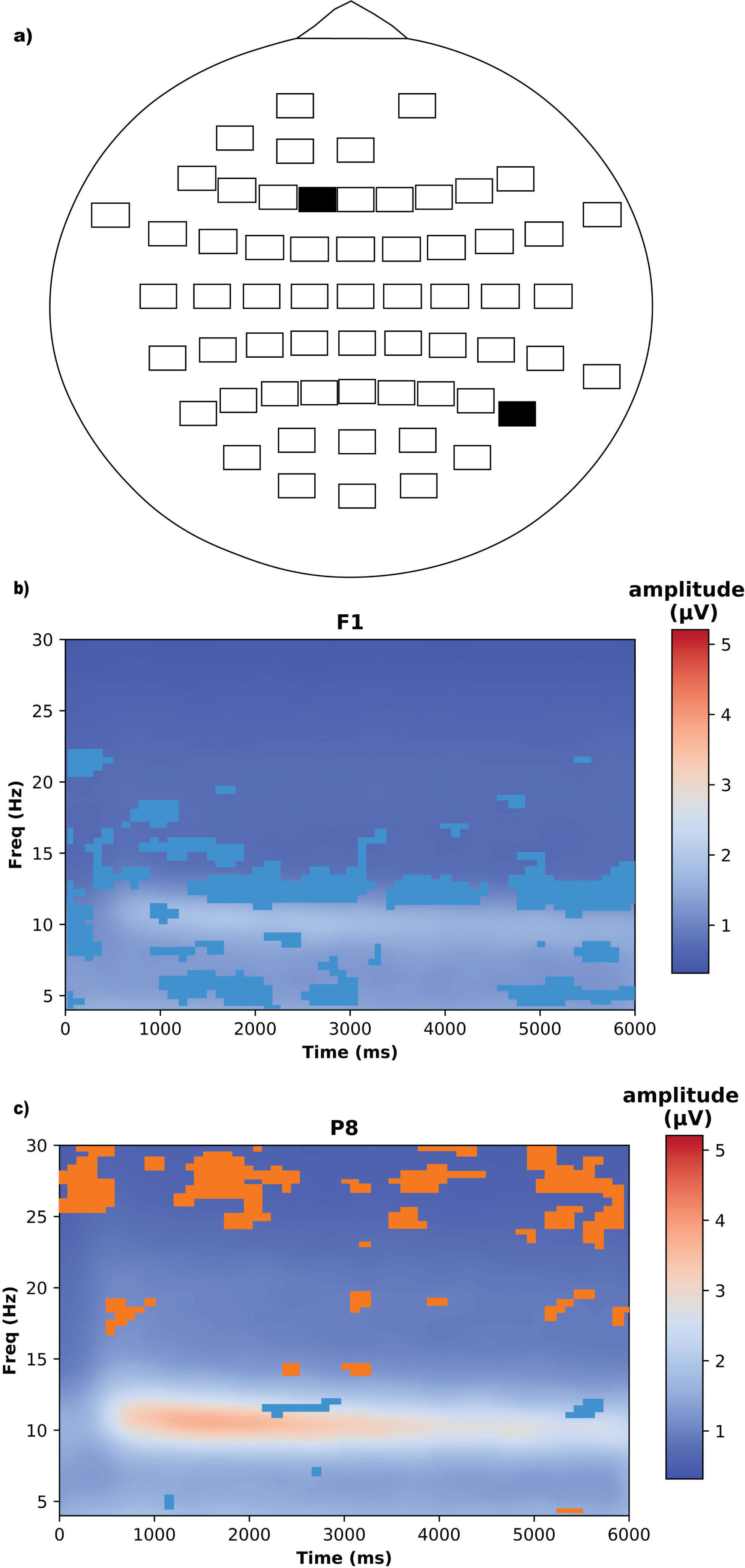
Comparison of Absolute Amplitude Delay Period Activity in Experiment 1 (Intact Scenes) Reveals Different Activity Patterns Between Rehearsal and Suppression. Select absolute amplitude plots in the left frontal and right parietal regions of the 6-sec delay period revealed 106 clusters of significant differences in activity (*p* < .05). The y-axis shows frequency (Hz); x-axis shows the time in sec. a) Montage of 61-scalp electrodes on head plot with two electrodes highlighted (colored black) F1 and P8. b) Selected electrode shown from the right parietal region (P8 electrode) displays orange clusters which represents the bins in frequency-time that are greater in amplitude for the rehearsal condition as compared with the suppression condition. Electrode selected based on results from previous research in our lab highlighting the right parieto-occipital region ^52^. c) Selected electrode from the left frontal region (F1 electrode) shows blue clusters which represents the bins in frequency-time that are greater in amplitude for the suppression condition as compared with the rehearsal condition.

Source-level analysis of the absolute amplitudes found 57 significant clusters across 13 of the 15 brain regions. The main source of activity for the rehearsal condition was central midline (CM_BR; 12 clusters, Time 100 to 6000 msec, Frequency 12-29 Hz), but overall changes were apparent over numerous brain regions. In the suppression condition, three regions were identified as sources of activity including the right temporal parietal region (TPR_BR; 8 clusters, Time 800 to 5400 msec, Frequency 10.5-17.5 Hz), the left temporal parietal region (TPL_BR; 8 clusters, Time 0 to 5900 msec, Frequency 7-12.5 Hz), and the right anterior temporal region (TAR_BR; 11 clusters, Time 100 to 5800 msec, Frequency 15-30 Hz) regions. The TPL_BR and TPR_BR amplitude differences are more concentrated in the upper alpha and lower beta ranges, whereas the TAR_BR activity was mainly in the beta range.

### Experiment 2

#### Behavioral

When task difficulty increased, there was both a significant short-term advantage of rehearsal (**Figure 2c and d**) compared to suppression (.85 vs. .78 proportion correct, *t*_(19)_ = 7.93, *p* < .001, *d* = 1.77) as well as a long-term advantage during rehearsal compared to suppression (.71 vs. .62 proportion correct, *t*_(19)_ = 4.58, *p* < .001, *d* = 1.02).

#### EEG

It was predicted that rehearsal and suppression would produce similar EEG delay period activity as in Experiment 1 since participants would be engaging in the same maintenance strategy, which should engage similar brain regions. Change in sensor-level amplitude over time (temporal spectral analysis) confirms a transient pattern of delay activity during rehearsal (Figure 3b).

Sensor-level temporal spectral analysis suggested transient changes, similar to the delay activity pattern observed in Experiment 1. Activity was increased and synchronous in the early delay period and became desynchronous later in the delay. Comparison between the rehearsal and suppression conditions using temporal spectral analysis revealed three clusters of significantly different activity (**Supplementary Table 3**). An early (0-1400 msec) and a late (3900-6000 msec) cluster represents alpha and beta ERD greater for the rehearsal condition as compared with suppression. The opposite pattern of TSA was identified in the middle of the delay period (1100-4600 msec) in which ERS was greater in the alpha and beta regions for the rehearsal condition as compared with suppression. Activity during rehearsal was also significantly correlated with WM performance (**Figure 5a-c**; Cluster 1: blue-negative correlation, cluster value = -38801.2, *p* = .005, Cluster 2: orange-positive correlation, Cluster value = 20445.5, *p* = .065), but not performance on the recognition task (**Figure 5d**; *p* = .62). Early ERS of alpha and beta activity was associated with better performance, and later ERD of the same frequency bands was associated with better performance on the WM task.

**Figure 5.**
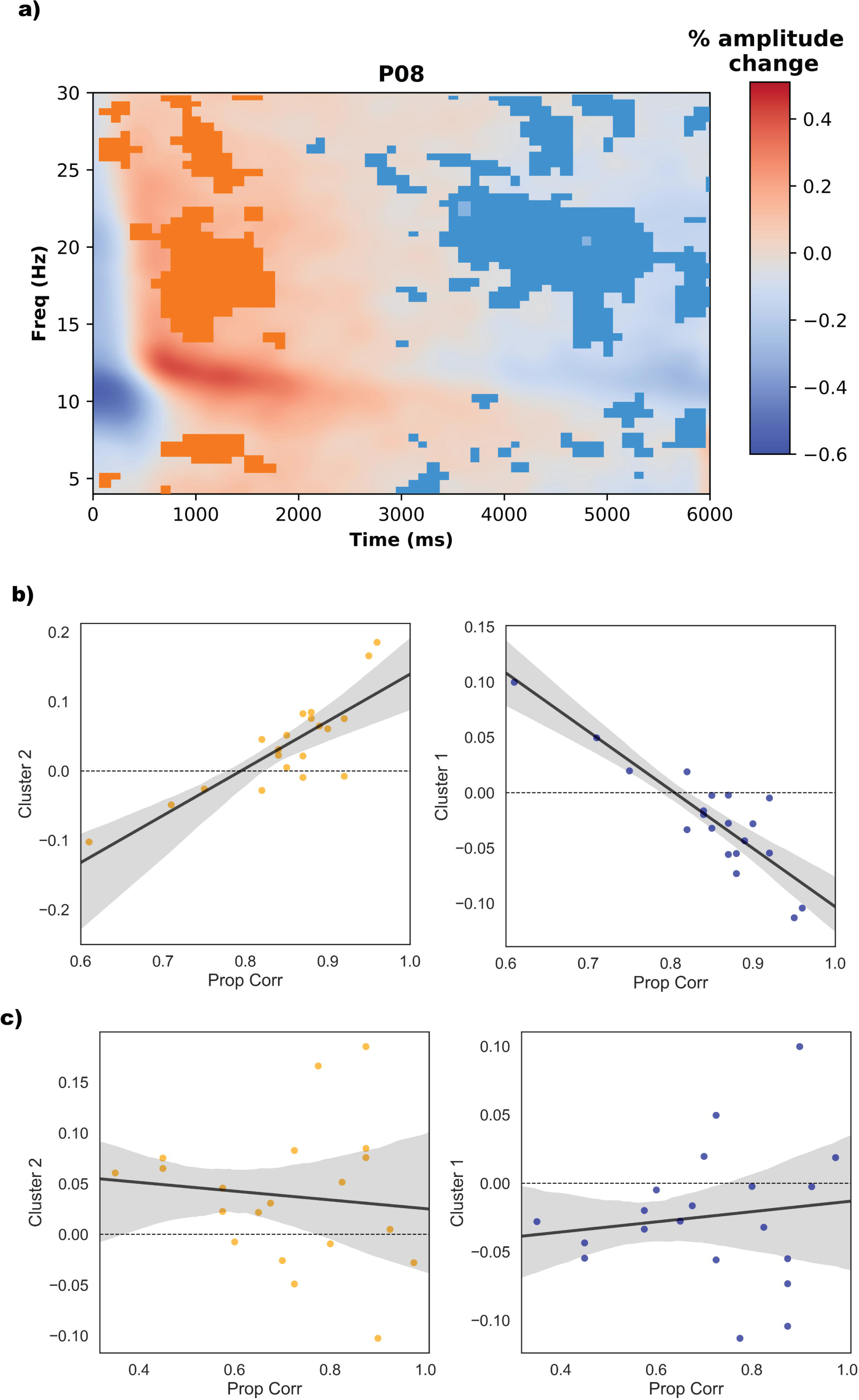
Delay Period Activity Time Frequency Analysis in Experiment 2 (Phase-Scrambled Scenes) Reveals Correlations with Performance. a) The y-axis is frequency (Hz); x-axis is the time in sec. Time Frequency Analysis correlation plot for the PO8 electrode during the 6-sec delay period in a whole window analysis. Significant clusters are represented by a mask of orange and blue. The first cluster (b, orange, latency: 0 to 2400 msec and frequency: 4 to 30 Hz) represents a positive correlation between activity during the delay period and performance on the working memory task from the rehearsal condition (Cluster value = 20445.5, *p* = .065) and the second cluster (c, blue, latency: 1400 to 4600 msec and frequency: 4 to 30 Hz) represents a negative correlation with performance (cluster value = -38801.2, *p* = .005). Performance on the recognition task was not significantly correlated with delay activity (d, *p* = .62).

Sensor-level analysis of the absolute amplitude between the two conditions (n = 20 subjects) with corrections for multiple comparisons revealed 15 significant clusters (**Supplementary Table 2**, *p* < .05). Greater amplitude was observed in the upper alpha and beta ranges across a majority of the sensors for the rehearsal condition as compared with the suppression condition, specifically in the bilateral temporal, occipital and parietal regions, and left frontal region (**Supplementary Figure 2**). The pattern of delay activity appeared to be sustained until about 3000 msec and then decreased in amplitude until the end of the delay period, specifically in right parietal and occipital regions (**Figure 6b** – P08 – orange cluster). There was also a brief early period (Time 0 to 1800 msec) in which greater amplitude was observed in the suppression condition in the theta, alpha, and beta range (blue cluster).

**Figure 6.**
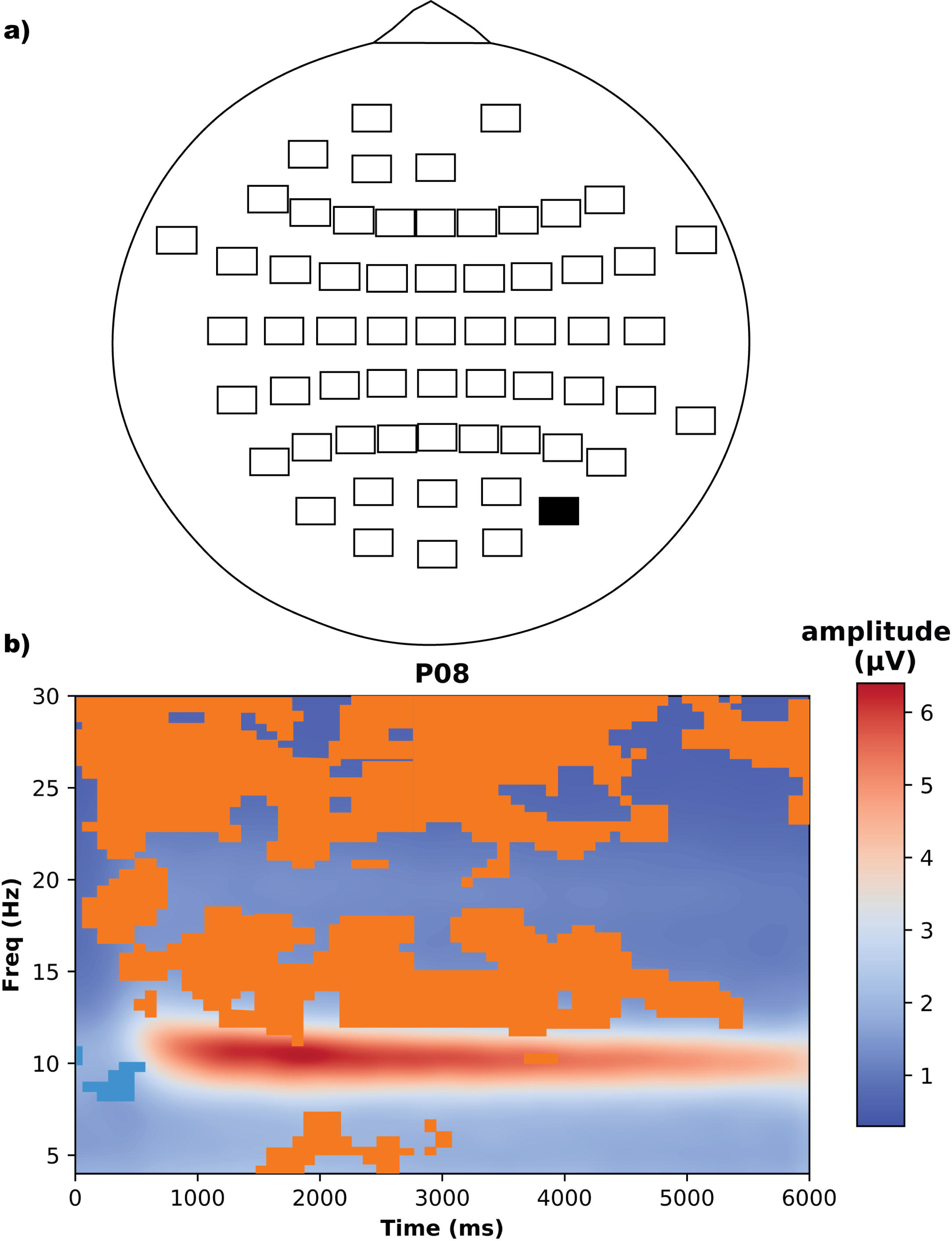
Comparison of Absolute Amplitude Delay Period Activity in Experiment 2 (Phase-Scrambled Scenes) Reveals Greater Activity for Rehearsal. The y-axis is frequency (Hz); x-axis is the time in sec. a) Montage of 61-scalp electrodes on head plot with one electrode highlighted (colored black) P08. b) Absolute amplitude plot for the PO8 electrode during the 6-sec delay period. PO8 was selected from all electrodes in the head map because it has the broadest clusters present of the other electrodes. The orange represents the clusters in frequency-time that are greater in amplitude for the rehearsal condition as compared with the suppression condition (15 significant clusters across all electrodes, p < 0.05). These cluster details are described in *Supplementary Table 3*.

Source-level analysis of absolute amplitudes revealed 67 significant clusters across 11 of the 15 brain regions (**Supplementary Figure 3**). Of the significant clusters, a focus of activity in the rehearsal condition was observed in the left anterior temporal region (TAL_BR; **Figure 7b**, 10 clusters, Time 0 to 6000 msec, Frequency 4-30 Hz), left parietal (PL_BR; 10 clusters, Time 0 to 6000 msec, Frequency 6-30 Hz), and midline parietal region (PM_BR; 6 clusters, Time 0 to 6000 msec, Frequency 20-30 Hz) regions. For the suppression condition, an early focal point of activity (Time 0 to 14000 msec) in the alpha range (Frequency 7-13.5 Hz) was found in the right and left parietal, parietal midline, and occipito-polar midline regions; a later focal point in the theta range (4-8 Hz) was found in the same brain regions as well as in the right frontal region (Time 3800 to 6000 msec). **Cross-Study Comparisons**

**Figure 7.**
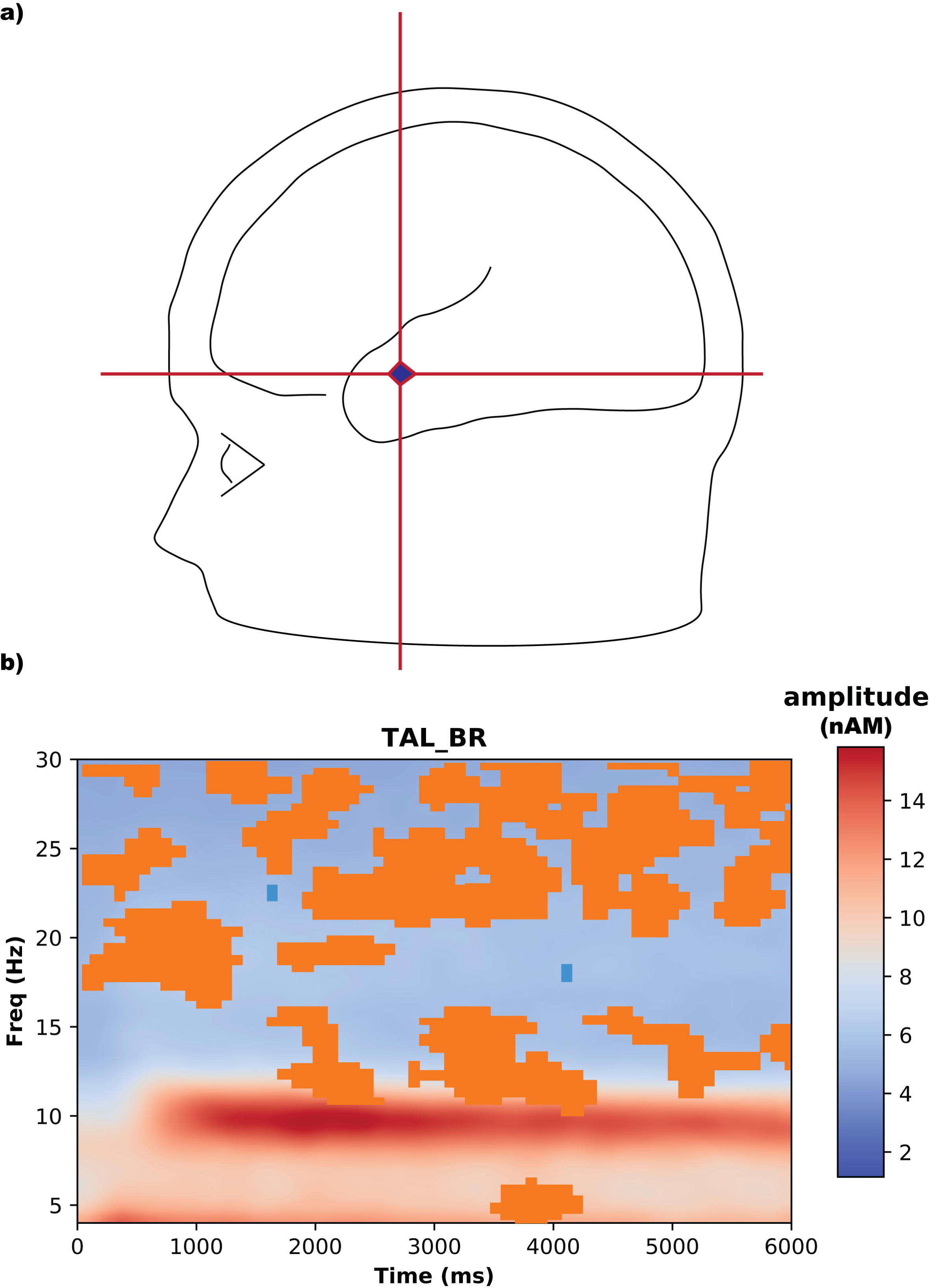
Delay Period Activity Brain Region Analysis in Experiment 2 (Scrambled Scenes) Reveals a Left Anterior Temporal Region during Maintenance Rehearsal. a) 3-D head with the left anterior temporal region (TAL_BR) highlighted. b) Absolute amplitude plot for the TAL_BR region during the 6-sec delay period (10 clusters, Time 0 to 6000 msec, Frequency 4-30 Hz). The y-axis is frequency (Hz); x-axis is the time in sec. Orange clusters represent rehearsal delay activity greater than suppression and blue clusters represent suppression delay activity greater than rehearsal. If focal activity is not visible, it suggests that the underlying sources of brain activity are more widespread. A clear focal point of activity for the rehearsal condition is found in the left anterior temporal region (TAL_BR) and the left- and middle-posterior regions (PL_BR, and PM_BR respectively see *Supplementary Figure 3*) which suggests that these regions are the source of delay activity for articulatory rehearsal.

Correlations between difference in absolute amplitude and difference in performance were run to see how changes in amplitude relate to changes in performance for Experiment 1 and Experiment 2. For Experiment 1, performance was not significantly different between conditions (**Supplementary Figure 4**). The differences in performance were not significantly correlated to differences in amplitude for any region or frequency range. In Experiment 2 however, Significant correlations between difference in absolute amplitude and difference in performance were found exclusively in the TAL_BR region. **Figure 8** shows the correlations for each region identified as a potential source of rehearsal by frequency. Increases in both Theta (*r* = 0.48, *p* = 0.034) and Alpha (*r* = 0.52, *p* = 0.019) oscillations are related to improved performance when rehearsal is employed. Importantly, the significant correlations for the TAL_BR region suggest that increases in theta and alpha band activity are related to improved short-term performance, when rehearsal is employed with images that lack semantic content.

**Figure 8.**
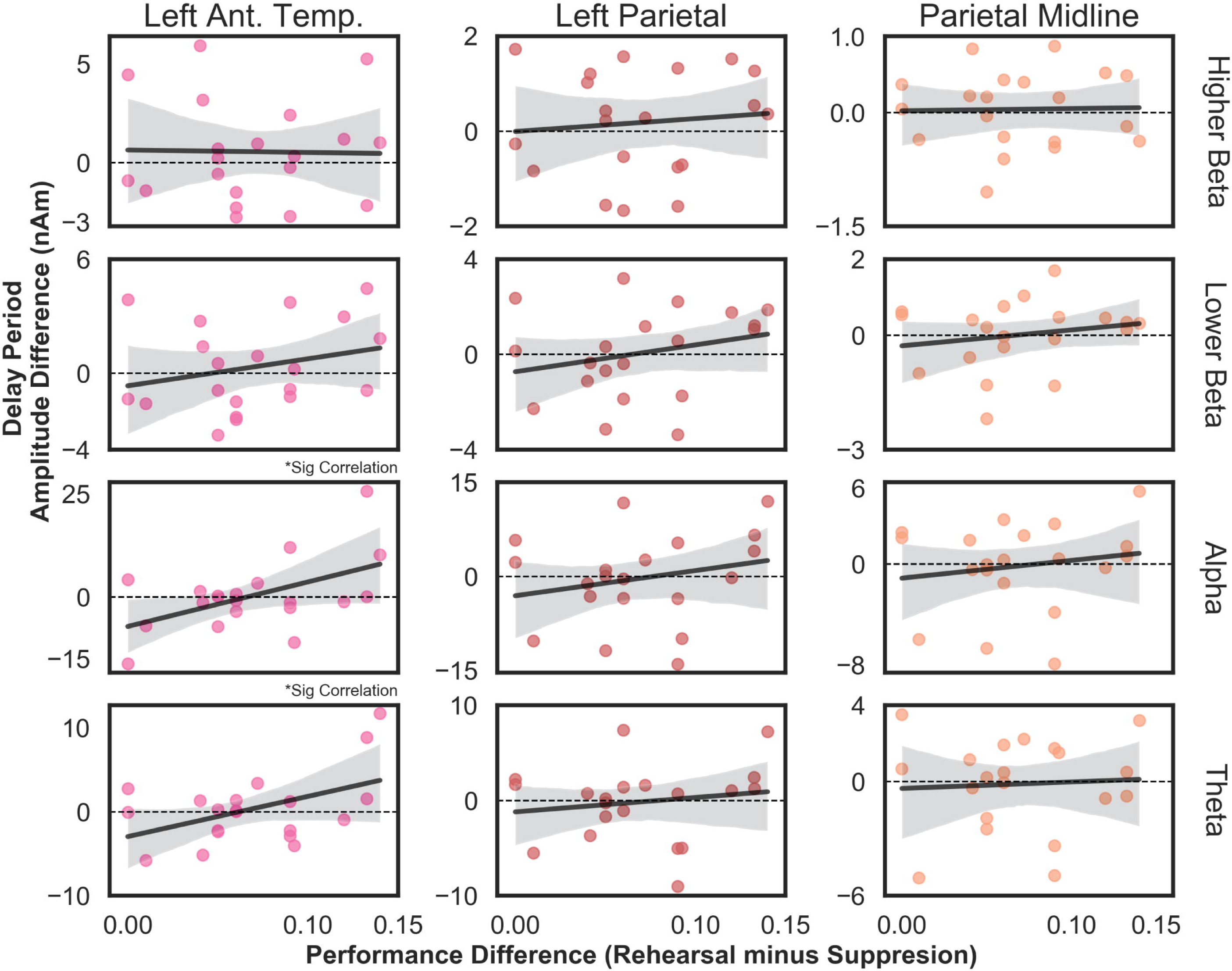
Correlations of Absolute Amplitude with Performance in Experiment 2 (Phase-Scrambled Scenes). The y-axis frequency difference (Amplitude for Rehearsal condition minus Amplitude for Suppression Condition). The x-axis performance difference (Proportion Correction for Rehearsal condition minus Proportion Correction for Suppression Condition). The frequency difference was calculated by averaging the absolute amplitude within a given frequency range (i.e. Theta (4-7 Hz), Alpha (8-13 Hz), Lower Beta (13-20 Hz), and Higher Beta (20-30 Hz) across the entire delay period. The correlations between difference in frequency and difference in performance are presented as a matrix for Experiment 1, with the TAL_BR region on the left (pink dots represent individual subjects), PL_BR region in the middle (red dots represent individual subjects), and PM_BR region on the right (orange dots represent individual subjects). These regions reflect the regions described in *Supplementary Figures 5-7*. These correlations reveal significant relationships in the Alpha and Theta frequency ranges for the TAL_BR region. Increases in both Theta (r = 0.48, p = 0.034) and Alpha (r = 0.52, p = 0.019) oscillations are related to improved performance when rehearsal is employed.

An exploratory comparison between results of Experiment 1 and Experiment 2 is included in the *Supplementary Results*.

## Discussion

The role that rehearsal plays in modulating delay activity remains an open research question. Experiments 1 and 2 examined how controlling for maintenance strategy impacted the delay activity observed during a visual WM task. A behavioral advantage of rehearsal was observed only when the stimuli lacked semantic content (i.e. phase-scrambled scenes). Examination of changes in delay activity over time during rehearsal with intact scenes (Experiment 1) and phase-scrambled scenes (Experiment 2) failed to reveal a sustained pattern of delay activity. Rather a transient pattern of delay activity with an early synchronous pattern in the alpha and beta ranges during the first half of the delay period followed by a desynchronous pattern of activity in the same ranges until the probe choice were observed. Additionally, early synchronous alpha and beta range activity and greater desynchronous activity in the same ranges was related to better short-term performance for the rehearsal condition with phase-scrambled scenes.

Examination of absolute amplitude during rehearsal with intact scenes (Experiment 1) and separately with phase-scrambled scenes (Experiment 2) revealed differences in all observed frequencies ranges when compared with suppression. Additionally, the TAL_BR, PL_BR, and PM_BR regions were identified as a source of delay activity during rehearsal. In fact, improved performance on rehearsal (as compared with suppression of rehearsal) in the former region (TAL_BR) was related to increases in the alpha and theta frequency bands. Interestingly, this was only the case when the stimuli lack-semantic content, suggesting that increases and alpha and theta are important when maintenance requires increased effort (i.e. automatic vs. non-automatic labeling).

In Experiment 1, generating a meaningful label for novel scenes and rehearsing them during the delay period provided neither a short- or long-term behavioral advantage. Comparison of sensor-level delay activity revealed a distinct pattern of greater amplitude in the beta range in the right parietal and centromedial regions for the rehearsal compared to suppression. This activity was not correlated with either short-nor long-term memory. Source-level analysis of EEG activity during rehearsal did not reveal any clear sources of activity. Additionally, comparison of change in amplitude over time did not reveal differences between the conditions. It did, however, reveal a transient pattern of change in amplitude in the alpha and beta ranges throughout the delay, with an early synchronous pattern followed by a late desynchronous pattern of activity. The lack of neural difference between change in amplitude over time between the suppression and the rehearsal condition, suggests that the same transient pattern of activity supported the near-ceiling performance in each condition.

When task difficulty increased in Experiment 2 using phase-scrambled scenes there was both a significant short-term as well as a long-term advantage with labeling and rehearsal. Sensor-level EEG activity during rehearsal showed greater amplitude in the upper alpha and beta ranges, specifically in the bilateral temporal, occipital and parietal regions and a left frontal region. Source-level analysis revealed a focus of activity during the rehearsal condition in the left anterior temporal, left parietal and parietal midline regions. Examination of the change in amplitude over time confirmed the transient pattern of delay activity that was observed in Experiment 1. Additionally, sensor-level temporal spectral analysis for rehearsal was significantly different from suppression and was also correlated with short-term memory with better performance associated with greater enhanced alpha and beta activity early in the delay period and greater suppression of these oscillations later in the delay period. The significant difference in neural activity between the suppression and the rehearsal condition suggests that increases in the alpha and beta ranges (early in the delay) and decreases in the same ranges (later in the delay) are modulated by the use of rehearsal and the amount of effort associated with recoding and rehearsing.

The role of rehearsal in supporting visual memory remains unclear, especially whether or not rehearsal benefits complex visual stimuli. Experiment 1 used intact, novel outdoor scenes that contained semantic information and were easy to generate a descriptive label for (e.g., a beach). In the rehearsal condition, label generation provided a dual means of encoding ^4,13,14^, one in the visual and one in the verbal domains, which was intended to benefit memory. However, there was no difference in performance on the short- or long-term memory task with intact scenes, which suggests that complex scenes do not benefit from this type of maintenance strategy. Complex scenes automatically trigger stored semantic associations ^12^; as a result, the association provided automatic deeper encoding ^11^ for both conditions. It has also been established that humans can remember thousands of images after only seeing the images for a brief time ^29-31^. This ability is termed the picture superiority effect ^32^ and may also account for the fact that performance for images was near ceiling regardless of maintenance strategy.

The benefit of a recoding a visually presented stimulus depends on whether semantic associations are automatically accessed without labeling ^5,33^. In Experiment 1 automatic semantic associations occurred; thus, the addition of rehearsing with a generated label offered no more benefit than accessing those stored associations ^12,14,26^. Although some stimuli such as spatial layouts may not benefit from recoding ^25^, some complex stimuli may benefit from this processes. In experiment 2, phase-scrambled images that lacked semantic content were used and these images benefited from recoding and rehearsal. This suggests that the benefit of rehearsal depends on the ability to associate semantic meaning; therefore, when participants generated a label and rehearsed throughout the delay period, they engaged in deeper encoding and elaborative rehearsal ^4,11,12,34,35^ improving short-term memory for these stimuli as well as long-term memory. These findings are consistent with the idea that generating a label and rehearsing is only beneficial to visual stimuli when semantic information is not automatically accessed ^14^ because it may direct the subjects’ attention to salient features ^5^ to support memory.

Delay activity during a working memory task is often associated with the engagement of either the prefrontal cortex or the posterior parietal regions but has been reported in studies that often fail to control for maintenance strategy. When intact scenes served as stimuli in Experiment 1, we observed greater activity in the left temporal and bilateral central regions during rehearsal, which suggests the engagement of the phonological loop ^22,36^. Suppression of rehearsal resulted in the engagement of more frontal regions suggesting reliance on attentional mechanisms ^37^ as well as greater mental effort ^38^. Source analysis revealed diffuse activity for both rehearsal and suppression conditions, which suggests no underlying focus of delay activity.

In Experiment 2, activity was greater in the upper alpha and beta ranges for the rehearsal condition throughout the delay period, specifically in the bilateral temporal, occipital and parietal regions and left frontal region, as compared with suppression. The brain regions analysis also revealed a focal source for the rehearsal condition in the left anterior temporal and parietal regions. Greater activation in the left temporal and frontal regions suggests the engagement of the phonological loop ^36,39,40^ and has also been implicated in the maintenance of non-spatial visual memory ^40^. Importantly, the anterior temporal lobe is associated with learning, memory as well as language. Bilateral temporal cortex resection in patients with epilepsy has been associated with reduction in short-term memory abilities, especially those involving semantic content ^41^. Left lateralized resection, as compared with right, has been shown to have greater impacts on learning of verbal information, short-term recall, and a greater rate of forgetting ^42,43^.

Additionally, the increased delay activity in frontal regions could represent the engagement of the central executive to support non-automatic recoding of difficult-to-label images, requiring increased attentional demand during rehearsal ^10^. Although, the phase-scrambled scenes contain the same visual features as regular scenes (i.e., color and spatial frequency), they lack the automatic semantic associations. The easy-to-label images used in Experiment 1, on the other hand, had an automatic semantic association and a verbal label ^10,12^. Thus, the generation of a label in Experiment 2 was more effortful (i.e., reliance on features ^6^) both during the recoding process and rehearsal and did not automatically produce a verbal label.

Engagement of occipito-temporal and parietal regions are associated with both visual working memory and attentional selection ^15,44^. Parietal lobe involvement in maintenance has been linked to attentional selection ^14,40^ specifically with regards to novelty ^44^ and may also play a role in the integration of features in complex objects ^15^. Although, activation in posterior parietal cortex has also been associated with attention, it likely does not exclusively reflect attentional processes ^24^. If recoding with semantic associations requires increased attention, as in the case of a more effortful association between a stimulus and stored semantic knowledge, the central executive is likely to play an important role as reflected by increased activation of frontal networks. If attentional resources are not required, as in the case of an automatic association such as in Experiment 1, the central executive is less likely to play a role ^26^. The results of these experiments confirm that controlling for maintenance strategy differentially recruits the prefrontal and parietal regions and modulates delay activity ^17^.

Elucidating how different patterns of delay activity contribute to WM is a current research focus ^17^. While it has long been established that sustained activity observed during maintenance represents both maintenance of encoded information and focusing of attention inward, recent research has suggested that delay activity is more complex ^17,28,45^. For example, only information in the focus of attention may be reflected in delay activity, while items outside the focus of attention may actually be represented by activity silent mechanisms ^45,46^.

Examination of the change in amplitude over time in both Experiments 1 and 2 suggests that when controlling for maintenance strategy, the pattern of delay activity is more transient. There is an early period of increased, synchronous activity (until approximately 3000 msec) as has been previously reported in the literature ^47-50^. The observed activity patterns were in the alpha and beta ranges, frequency ranges that are critical for maintenance of visual information. Alpha enhancement likely reflects inhibition of information that could interfere with maintenance, while beta activity is related to controlling the information that is being actively maintained ^51^. The early delay activity was followed by a period of desynchronous activity for the remainder of the delay which may suggest preparation for the upcoming response ^52^. This pattern of activity is consistent with recent reports that maintenance is not necessarily supported by persistent delay activity in prefrontal regions ^17,28^; instead, delay activity may reflect more complex processes in cortex and subcortical regions. Alternatively, previous reports of sustained delay activity could reflect a maintenance period in which participants did not utilize a particular strategy, rather, they focused their attention inward until they were required to produce a response ^27^.

We argue that greater effort is likely required to maintain difficult-to-name complex visual stimuli (i.e. phase-scrambled scenes), as compared with intact scenes, because the labeling and rehearsal process is not automatic as it is with intact scenes. Although, this was not evident in the experiment-level comparison between absolute amplitude, it can be argued that it is evident in the correlations between absolute amplitude and performance. In Experiment 2, increases in theta and alpha band activity were associated with increased performance with the application of rehearsal. Interestingly, performance also improved for some subjects that showed a decrease in theta and alpha band activity. The theta and alpha rhythms have long been implicated in working memory maintenance ^48,50,53^. Theta has been found to play an important role in working memory maintenance, specifically in frontal regions ^53^, and may be related to focusing attentional resources on the to-be-remembered stimuli ^54^. More recently, theta has been associated with language processing ^55^, including language comprehension ^56^, language generation ^57^, and semantic associations ^58^. More specifically, studies of semantic processing using sentence violation paradigms have found increases in theta power in the hippocampus, parahippocampus as well as the language-associated neocortical areas ^58^ specifically with processing semantic meaning. An opposite pattern of decreases in theta power have also been reported in the hippocampus, specifically during generation of the language ^57^.

Although the Experiment 2 findings of both increases and decreases in theta are related to improved performance is contradictory, the success of both groups of subjects may reflect individual application of the same technique. The cognitive operations involved generating a label to describe the picture followed by rehearsal of the label. Subjects were encouraged to use meaningful labels, as in Experiment 1, but required more effort in the generation as the association was not automatic. For some participants, drawing on long-term semantic associations provided a behavioral advantage when rehearsal was applied, which was evident in increased theta band activity ^58^. Whereas, for others more shallow, less meaningful labels were applied; therefore, the rehearsal period may simply reflect the generation of words that are rehearsed, as evidenced by a decrease in theta band activity ^57^. This method was equally as effective in the short-term, likely because of the low working memory load and short period over which maintenance occurred.

One limitation of the current study is that participants engaged in covert rather than overt rehearsal and suppression. Covert rehearsal was used to reduce the amount of noise introduced into the EEG signals. Task compliance was therefore based on participant confirmation during the recognition task (i.e., they reported their generated labels). It is possible that participants did not engage in suppression; consequently, the failure to produce the intended behavioral outcome in Experiment 1 could be explained by this. While the intended behavioral outcome was found in Experiment 2, it is again possible that participants did not engage in suppression. Future experiments that rely on covert maintenance mechanisms should include a response-based behavioral confirmation such as an immediate reporting of generated labels or the number of times the suppression word is repeated. Future studies should also examine the type of label generated to see if the depth of label (i.e., deeper descriptive label like “sunset” vs. shallower label like “white and blank lines”) or the verbalizability of the stimulus ^26^ impacts subsequent memory.

## Conclusions

The results of these experiments provide behavioral evidence that rehearsal impacts short-term memory for some, but not all visual stimuli. Rehearsal also modulates the neural pattern of delay period activity as represented by the absolute amplitude of the delay period signal, especially in the left anterior temporal region. Changes in theta and alpha bands during the delay period correlate with working memory performance but only when maintaining labeled associations for stimuli that lack semantic content.

## Materials and Methods

### Participants

The study was approved by the Institutional Review Board of the City University of New York Human Research Protection Program (CUNY IRB). A total of 54 participants provided informed consent and completed the study. The study was carried out following the CUNY IRB guidelines and regulations. Participants were compensated with either 15 dollars or one extra course credit per hour of participation. The behavioral task with EEG recordings took approximately 2 hours to complete.

#### Experiment 1

One participant was excluded from Experiment 1 of the study for failure to follow instructions resulting in a final sample in Experiment 1 of 29 participants (age = 25.4 (8.1) years, 14 females). For EEG analysis a total of 6 participants were excluded: 5 participants for noisy or unusable EEG recordings and 1 participant for failing to follow instructions resulting in a final sample for Experiment 1 consisting of 24 participants (age = 25.8 (8.6) years, range 18-56, 11 females).

#### Experiment 2

Experiment 2 was a replication of Experiment 1 with phase-scrambled stimuli to increase task difficulty by removing semantic content. A complex visual stimulus that has not been explored to our knowledge is a phase-scrambled scene which is difficult to recode, specifically with one-word representations, and is often described based on basic features (i.e. colors or line orientation). The difficult-to-describe nature of this stimulus makes it suitable for studying visual memory because it is not likely to automatically be recoded nor trigger a deeper semantic representation ^4,26^. Therefore, it is well suited to assess benefits of maintenance strategies (i.e. articulatory rehearsal) for storing and remembering complex visual information.

Four participants were excluded because of computer malfunction preventing recording of behavioral responses resulting in a final sample of 20 participants (age = 24.8 (9.5) years, range 18-56, 12 females).

### Experimental Design and Statistical Analyses

#### Task

Participants completed a modified version of a Sternberg WM Task ^59^. The task consisted of two WM tasks (**Figure 1a and b**) and a delayed recognition task. During the WM task, participants completed a low load (2 images) WM task consisting of an encoding phase, delay period, probe choice response, and a phase-scrambled image that indicated the end of the trial. During presentation of the images, participants were instructed to generate a verbal label (i.e. descriptive label like “beach”), and during the delay period they were instructed to rehearse covertly (i.e. using their inner voice) the label throughout the entire delay period (rehearsal) or were prevented from actively rehearsing (suppression). Participants completed both the rehearsal and suppression conditions in a randomized order. The delayed recognition task was administered ten minutes later.

For suppression, participants were discouraged from generating a verbal label and instructed to repeat the word “the” throughout the delay period ^60,61^. Suppression was used to block verbal rehearsal by engaging the articulatory process of the phonological loop with an unrelated word. This suppression task uses repetition of a non-task related word such as “the” over and over again ^60-62^. This classic suppression task was chosen over an alternative task such as random number generation or a math problem because these alternative tasks engage attentional processes. The goal of suppression is to block rehearsal without a demand on attention, which has been demonstrated to have a longer term effect on memory tasks ^62,63^.

#### Stimuli

In Experiment 1, stimuli consisted of high-resolution, color outdoor scenes, which did not contain any faces or words. Scenes were randomly selected from the SUN database ^64^ and were resized to 800 by 600 pixels.

Experiment 2 employed the same design as Experiment 1 with Fourier phase-scrambled (Matlab, v2016) versions of the scenes used in Experiment 1 (**Figure 1c and d**). Importantly, the scenes contained the same colors and spatial frequencies as the scenes used in Experiment 1 but lacked semantic content. It was more challenging to generate labels because phase-scrambling removes all semantic content.

#### Behavioral Analysis

Behavioral data were processed in Python 3.0, and plots were created using Seaborn 0.9.0 with custom scripts. Statistical analysis was conducted in Jasp v0.9.0.1 using paired-samples t-tests to compare behavioral accuracy between conditions (rehearsal vs. suppression) on the WM and recognition tasks and Pearson correlations between behavior and delay activity.

#### EEG Acquisition, Processing and Plots

Continuous 64-channel EEG was collected at 1000 Hz using an active electrode system (actiCHamp, Brain Products). A 10-20 montage was used with a left mastoid reference (TP9). Raw EEG data were processed in BESA Research v6.1. Data were visually inspected, eye blinks were corrected with a pattern-matching algorithm ^65,66^, and muscle artifacts were removed. Participants were included if they had more than 50 trials that survived the artifact scan (*see Supplementary Methods*).

Time-frequency analyses (TFA; absolute amplitude and temporal spectral analysis) were conducted on artifact-corrected delay periods (0 to 6000 ms) and bandpass-filtered between 4 and 30 Hz. Complex demodulation of the recorded EEG signals for each trial was carried out in BESA Research v6.1 ^67,68^. A detailed description of the demodulation can be found in Ellmore, et al. ^52^. Briefly, the timeframe (t) was set as the entire delay period (0 to 6000 msecs) and the baseline was set to the same period ^52^. The finite impulse response (FIR) filter was applied in latencies of 100 msecs and 0.5 Hz steps. First, the raw amplitude of brain activity within each frequency and latency bin was generated for condition (termed Time Frequency Analysis or TFA). Then, the amplitude within in frequency bin across time was compared to the baseline amplitude and averaged over trials (termed Temporal Spectral Analysis or TSA). The TSA is:

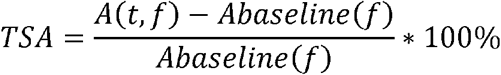

A(t,f) = amplitude during the timeframe of interest and frequency A_baseline_ (f) = mean amplitude in the frequency band during the baseline period

Temporal Spectral Analysis is expressed as a percent or change in amplitude, the resulting TSA is a value that is either + or -. A positive change represents enhancement of activity (or synchronization) and a negative change represents a suppression of activity (or desynchronization) relative to the event of interest, the delay period ^69,70^.

A brain source montage was applied to TFA to distinguish between broad, potentially overlapping sources (i.e. frontal, central, parietal, etc.). The brain region montage consists of 15 discrete regions and source-level analysis calculates weighted combinations of the recorded scalp electrodes in order to reduce the amount of overlap^39,67^. This source-level analysis provides a powerful way to reduce the spatial overlap of the sensor-level findings ^52^.

A Brain Vision Matlab script for reading BESA-generated files into other packages (https://www.besa.de/downloads/matlab/) was adapted to read in the TFA files generated into Python. The TFA average plots were recreated using custom scripts. The significant clusters from permutation tests that were generated in BESA Statistics, were retraced in Adobe Illustrator v2019, and overlaid on the python-generated TFA average plots. Average TFA by frequency were calculated using custom Python scripts. The frequency band differences by band were calculated by averaging the absolute amplitude within a given frequency band across the entire delay period (0-6000 msec) for rehearsal and subtracting the corresponding value for the suppression condition. The frequency (f) ranges under consideration were 4 to 30 Hz: theta (4-8 Hz), alpha (8-13 Hz), and lower and upper beta (13-20 and 20-30 Hz, respectively).

#### EEG Statistical Analysis

Scalp-level TFA and source-level TFA were compared using paired-samples t-tests with corrections for multiple comparisons using non-parametric cluster permutation testing (N = 1,000 permutations) ^71^ in BESA Statistics v2.0. Correlations were run between TFA and performance with corrections for multiple comparisons. In order to account for multiple comparisons across frequency, time, and sensor space, non-parametric permutation t-tests or correlations were run. The analyses result in a cluster value. For the t-test, the cluster value is a sum of t-values and for a correlation, the cluster value is a sum of the r-values for the data points in a group of adjacent bins (sensor (<4 cm distance), time (100 msec), frequency (.50 Hz) bins). A null distribution of summed t-values for the t-test analysis or r-values for the correlation analysis are generated from random clusters across subjects and across time-frequency bins. Significant clusters are summed t- or r-values within a time-frequency domain that exceed a specific threshold and are then compared to the random null distribution (8). Significant clusters were defined as p-value less than or equal to .05 and marginally significant was less than or equal to .01. Cluster permutation tests involve sensors that are <4 cm apart, which suggests that if a significant finding occurs across neighboring electrodes it is unlikely that the effect is due to chance. This type of non-parametric permutation test overcomes the Type I error that can occur with multiple comparison testing. Significant clusters are indicated with a mask, either blue or orange. An orange cluster represents greater amplitude for one condition versus another (always rehearsal as compared with suppression) for the TFA or event-related synchronization for the TSA. Blue represents a smaller amplitude (for rehearsal versus suppression) for the TFA or event-related desynchronization for the TSA.

## Supporting information

Supplemental Methods and Results

## Acknowledgements

We thank Ashley M. Arango for help with task development, data collection and EEG processing. We would also like to thank Ning Mei for help with coding and script development, and Farzana Antara, Bernard Gomes, and Jefferson Ortega for feedback on the manuscript.

## Availability of Data

The datasets generated during and/or analyzed during the current study are not publicly available because the approved IRB protocol does not include a provision for deposit in a public repository, but the de-identified data are available from the corresponding author TME on reasonable request.

## Author Contributions

C.R.P and T.M.E. conceived the experiment. C.R.P. and K.N contributed to task development, data collection, and data processing. C.R.P. carried out all data analysis and wrote the manuscript. K.N. reviewed and provided feedback on all versions of the manuscript. T.M.E. aided in interpreting the results and writing the manuscript as well as supervised all aspects of the experiment.

