## Supplemental Methods and Results for "Does rehearsal matter? Left anterior temporal alpha and theta band changes correlate with the beneficial effects of rehearsal on working memory"

Chelsea Reichert Plaska

Kenneth Ng

Timothy M. Ellmore

**Supplementary Methods**

**Task**

Participants completed a modified version of the Sternberg working memory task, separable (e.g. encoding, delay period, probe choice). The overall study design consisted of two working memory (WM) tasks (see **Fig. 1**; 100 trials per task) and a delayed old-new recognition task (150 trials) approximately 10-minutes later. For the WM tasks, participants were presented with a fixation cross (1 sec) that indicates the start of the trial, followed by 2 images in succession (2 sec each), a blank screen during the delay period (6 sec), a probe choice (2 sec), which is either one of the earlier presented images or a new image, and a phase-scrambled image (1 sec) that indicates the end of the trial. The long 6-sec delay-period is indicative of real-world scenarios for maintaining information and have been used in similar studies of memory ^1^. During this short 10-minute break, participants remained in the lab. For the working memory tasks, participants were given examples of verbal labels during a short 10-trial practice set as well as the rate at which they should rehearse or suppress, before beginning the task. The participant made probe choices using a RB-530 response pad (Cedrus Inc). If a probe matched one of the previously presented images from the encoding set, the participant pressed the green (right) button on the response pad. If the probe did not match the encoding set, the participant pressed the red (left) button.

The recognition task was a mix of any encoding image from either the rehearsal and suppression conditions (40 images from each condition), as well as new images (70 images), participants indicated if the images were old or new. During the recognition task, the participant indicated if the image was presented in either of the working memory conditions or if it was a new image. If they indicated that they saw the image in one of the earlier working memory conditions, they were asked to indicate if they remembered labeling the image and verbally stated the label that was used. The recognition task was a forced-choice task. Participants were given unlimited time to provide the label overtly. This confrontation of participants during the recognition task by the experimenter or a research assistant served as task-check to see if participants followed instructions. All participants, except one who was excluded from the analysis, confirmed that they followed instructions and indicated the condition in which they saw the old images (i.e., rehearsal vs suppression). Although the generated labels were reviewed during the recognition task to confirm task compliance, they were not systematically analyzed.

The stimuli were presented as 800 by 600 pixels on a gray background. Precise stimulus timings were recorded using a photosensor that was located on the computer monitor in the upper right corner. The photosensor detected the changes using a small black box presented on the gray background. The small box was present whenever a stimulus was present and was not visible to participants. The timings were then coded to represent the separate events (i.e. delay period, probe, etc.).

**EEG Acquisition and Processing**

Continuous 64-channel EEG was collected at a sampling rate of 1000 Hz using an active electrode system with actiCHamp system (Brain Products). Electrodes were reduced to an impedance of 25 kOhms or lower, per the manufacturer’s specifications. Channels with an impedance above 25 were interpolated. The reference electrode was the left mastoid (TP9) and was re-referenced offline to the common average reference. Data were visually inspected and muscle artifacts were identified and marked for removal. Eye blinks were corrected with a pattern-matching algorithm using principle component analysis ^2,3^. The method removes the variance associated with a blink, from each channel, using LOC and ROC channels (opposite deflection pattern > 100 μV). Artifact-corrected data was used for all analyses.

Participants were included if they had more than 50 delay periods that survived the artifact scan in both the rehearsal and suppression conditions. Trials were excluded based on BESA Research criteria for artifact rejection: trials were excluded for amplitude > 148 μV*, gradient > 75 μV, low-signal criteria > 0.1. In order to maximize the number of trials included, amplitudes (*) above 148 were manually reviewed and may have been included on individual trial basis. In Exp 1, the average number of delay period was 77 for suppression and 78 for rehearsal and in Exp 2, it was 80 for each condition, with no significant differences between the number of delay periods included (*p* > .05).

**Supplementary Results**

A formal comparison between brain activity in Experiment 1 and Experiment 2 was not planned. As a result, there were no a priori hypotheses regarding the comparison. The exploratory analysis between Experiments 1 and 2 was conducted using a between-subjects analysis of variance (ANOVA) with corrections for multiple comparisons using non-parametric permutation testing as described in the Supplementary results. In order to reduce the number of comparisons, the delay periods were averaged across trials, subjects, and then channels. A between-subjects ANOVA was conducted for absolute amplitude and TSA at the sensor-level, separately, treating each condition (Experiment 1-rehearsal, Experiment 1-suppression, Experiment 2-rehearsal, Experiment 2-suppression) as independent groups. For absolute amplitude, there were no significantly different clusters of brain activity between the groups. Similarly, for TSA there were no differences between the groups. These findings suggest that the same brain pattern activity was exhibited during rehearsal and suppression conditions, regardless of the type of image.

Further examination of the distribution of absolute amplitude values for each condition was conducted within each frequency range (Theta (4-7 Hz), Alpha (8-13 Hz), Lower Beta (13-20 Hz), and Higher Beta (20-30 Hz)) for selected regions found to be critical for rehearsal based on the Brain Regions Analysis (see Figure 8) were chosen: the Left Anterior Temporal Region (TAL_BR), the Left Parietal Region (PL_BR), and the Parietal Midline Region (PM_BR). The absolute amplitude values were calculated by averaging the entire delay period within a given frequency range for each subject, for the rehearsal and for the suppression conditions. For Experiment 2, increases occurred in about 50-60% of subjects across all frequency ranges. The overall distributions (i.e. violin plots and boxplots) for each condition and experiment reveal overlapping distributions within each frequency range for all three regions. **Supplementary Figure 5** shows the distributions for each experiment and condition for the TAL_BR for the Theta, Alpha, and Beta ranges (see **Supplementary Fig. 6 and 7** for PL_BR and PM_BR**)**. In both experiments, between 50-60% of subjects showed an increase in amplitude from the suppression condition to the rehearsal condition for all frequency bands, but the overall distributions for each conditions are overlapping, which provides for support for no neural difference between the experiments.

**Discussion and Limitations**

The exploratory analysis comparing the conditions (rehearsal and suppression) between Experiments 1 and 2 showed that overall pattern of delay activity was not significantly different between the experiments. Although, behaviorally one type of complex visual stimuli (i.e. phase-scrambled scenes) may benefit from rehearsal, this difference was not reflected in the cross-study comparison of delay activity. One explanation is that both experiments revealed a transient patterns of delay activity with early synchronous component and a late desynchronous component focused in the alpha and beta frequency ranges. These patterns of activity are essential to successful working memory maintenance and likely reflect different but complementary cognitive processes. Therefore, the fact that there was no neural difference between experiments suggest that the overall change in delay activity across an extended maintenance period is similar for different types of visual stimuli. Alternatively, the lack of neural difference between experiments may be the result of the analysis which involved different groups of subjects in each condition, as compared with the original analyses that were within-subject analyses. Therefore, this may be due to low power and uneven, small sample sizes, especially for Experiment 2. Future studies should compare delay activity with different task demands using the same subjects in each group to examine the interaction between rehearsal and stimulus type.

A major limitation of these analyses is that the participants in Experiment 1 were different participants from those in Experiment 2. Additionally, due to limitations in our statistical software, each condition (rehearsal and suppression) were treated independently; thus, there was no deliberate attempt to control for confounding variables (i.e. age, education level, etc.) or differences in performance. As described in the planned analyses, rehearsal and suppression were completed by the same participant for each experiment and the appropriate paired-samples analyses were conducted. Additionally, there was no attempt to separate participants based on their working memory capacity, into high and low working memory capacity groups. Delay activity may vary as a function of working memory capacity as there are individual differences in the ability to inhibit task-irrelevant information. This could potentially have explained the large variability in absolute amplitude the lower frequency ranges for some participants ^4^ and should be considered for future experiments.

**Supplementary Figure 1. Comparison of Absolute Amplitude Delay Period Activity in Experiment 1 Reveals Different Activity Patterns Between Rehearsal and Suppression.** Select absolute amplitude plots in the left frontal and right parietal regions of the 6-sec delay period revealed 106 clusters of significant differences in activity (*p* < .05). The y-axis shows frequency (Hz); x-axis shows the time in sec. Head plot of the overall pattern of absolute amplitude difference during the delay period for all sensors. Orange clusters represent rehearsal delay activity great than suppression and blue clusters represent suppression delay activity great than rehearsal. These cluster details are described in *Supplementary Table 1*.

**Supplementary Figure 2.** **Comparison of Absolute Amplitude Delay Period Activity in Experiment 2 Reveals Greater Activity for Rehearsal.** The y-axis is frequency (Hz); x-axis is the time in sec. Head plot of the overall pattern of absolute amplitude difference during the delay period for all sensors. These cluster details are described in *Supplementary Table 2*. Orange clusters represent rehearsal delay activity great than suppression and blue clusters represent suppression delay activity great than rehearsal. These cluster details are described in *Supplementary Table 3*.

**Supplementary Figure 3. Delay Period Activity Brain Region Analysis in Experiment 2 (Scrambled Scenes) Reveals a Left Anterior Temporal Source during Maintenance Rehearsal.** The y-axis is frequency (Hz); x-axis is the time in sec. Head plot of the overall pattern of absolute amplitude difference during the delay period for all sensors grouped by brain region. Orange clusters represent rehearsal delay activity greater than suppression and blue clusters represent suppression delay activity greater than rehearsal. If focal activity is not visible, it suggests that the underlying sources of brain activity are more widespread. A clear focal point of activity for the rehearsal condition is found in the left anterior temporal region (TAL_BR, see *Figure 7*) and the left parietal region (PL_BR) which suggests that these regions are the source of delay activity for articulatory rehearsal. No clear focal point of activity emerged for the suppression condition, which suggests that the source of activity is more diffuse.

**Supplementary Figure 4. Correlations of Absolute Amplitude Difference and Performance in Experiment 1 (Intact Scenes).** The y-axis frequency difference (Amplitude for Rehearsal condition minus Amplitude for Suppression Condition). The x-axis performance difference (Proportion Correction for Rehearsal condition minus Proportion Correction for Suppression Condition). The frequency difference was calculated by averaging the absolute amplitude within a given frequency range (i.e. Theta (4-7 Hz), Alpha (8-13 Hz), Lower Beta (13-20 Hz), and Higher Beta (20-30 Hz) across the entire delay period. The correlations between difference in frequency and difference in performance are presented as a matrix for Experiment 1, with the TAL_BR region on the left (pink dots represent individual subjects), PL_BR region in the middle (red dots represent individual subjects), and PM_BR region on the right (orange dots represent individual subjects). These regions reflect the regions described in *Supplementary Figures 5-7*. These correlations show that there was no relationship between performance and changes in absolute amplitude within the selected frequency ranges and regions.

**Supplementary Figure 5. Comparison of Absolute Amplitude During the Delay Activity for Rehearsal and Suppression for Experiment 1 (Intact Scenes) and Experiment 2 (Phase-Scrambled Scenes) in the Left Anterior Temporal Region suggest overlapping distributions.** Absolute Amplitude for the Left Anterior Temporal Region (TAL_BR) is shown with four plots for each experiment. Experiment 1 (Intact Scenes) is shown on the left and Experiment 2 (Phase-Scrambled Scenes) is shown on the right. The TAL_BR region was selected based on the findings presented in *Figure 8.* Each plot represents a frequency range: Theta (4-7 Hz), Alpha (8-13 Hz), Lower Beta (13-20 Hz), and Higher Beta (20-30 Hz). The absolute amplitudes that are plotted in each range represent the average within that range across the entire 6-second delay period. 1) The violin plot shows the distribution of amplitude across each condition; the cluster of similar amplitude values is reflected at the wider (bottom) portion of the violin and the thinner (top) portion with fewer amplitude values. The dashed lines reflect the quartiles of the overall distribution similar to the box plot. 2) The boxplot also shows the distribution of performance for each condition (i.e. the solid lines inside the box align with the dashed lines of the violin plot, representing the quartiles). In addition, the boxplot contains whiskers extend to the minimum and maximum score (1.5 times the median) for each condition and the diamonds reflect outliers (great than 1.5 times the median). Outliers were included in all analyses. 3) In between the violin plot and boxplot is a point plot, which reflects the mean of absolute amplitude, with error bars reflecting the 95% confidence intervals; the confidence intervals were generated with bootstrapping using 1,000 iterations. 4) Finally, the individual dots on the inner-most portion represent a single participants absolute amplitude for a condition (Experiment 1 *n = 24* and Experiment 2 *n* = 20). The gray line connects the dots that represents absolute amplitude value for the corresponding condition (i.e. suppression (left) and rehearsal (right)).

**Supplementary Figure 6. Comparison of Absolute Amplitude During the Delay Activity for Rehearsal and Suppression for Experiment 1 (Intact Scenes) and Experiment 2 (Phase-Scrambled Scenes) in the Left Parietal Region.** Absolute Amplitude for the Left Parietal Region (PL_BR) is shown with four plots for each experiment. Experiment 1 (Intact Scenes) is shown on the left and Experiment 2 (Phase-Scrambled Scenes) is shown on the right. The PL_BR region was selected based on the findings presented in *Figure 8.* Each plot represents a frequency range: Theta (4-7 Hz), Alpha (8-13 Hz), Lower Beta (13-20 Hz), and Higher Beta (20-30 Hz). The absolute amplitudes that are plotted in each range represent the average within that range across the entire 6-second delay period. 1) The violin plot shows the distribution of amplitude across each condition; the cluster of similar amplitude values is reflected at the wider (bottom) portion of the violin and the thinner (top) portion with fewer amplitude values. The dashed lines reflect the quartiles of the overall distribution similar to the box plot. 2) The boxplot also shows the distribution of performance for each condition (i.e. the solid lines inside the box align with the dashed lines of the violin plot, representing the quartiles). In addition, the boxplot contains whiskers extend to the minimum and maximum score (1.5 times the median) for each condition and the diamonds reflect outliers (great than 1.5 times the median). Outliers were included in all analyses. 3) In between the violin plot and boxplot is a point plot, which reflects the mean of absolute amplitude, with error bars reflecting the 95% confidence intervals; the confidence intervals were generated with bootstrapping using 1,000 iterations. 4) Finally, the individual dots on the inner-most portion represent a single participants absolute amplitude for a condition (Experiment 1 *n = 24* and Experiment 2 *n* = 20). The gray line connects the dots that represents absolute amplitude value for the corresponding condition (i.e. suppression (left) and rehearsal (right)).

**Supplementary Figure 7. Comparison of Absolute Amplitude During the Delay Activity for Rehearsal and Suppression for Experiment 1 (Intact Scenes) and Experiment 2 (Phase-Scrambled Scenes) in the Parietal Midline Region.** Absolute Amplitude for the Parietal Midline Region (PM_BR) is shown with four plots for each experiment. Experiment 1 (Intact Scenes) is shown on the left and Experiment 2 (Phase-Scrambled Scenes) is shown on the right. The PM_BR region was selected based on the findings presented in *Figure 8*. Each plot represents a frequency range: Theta (4-7 Hz), Alpha (8-13 Hz), Lower Beta (13-20 Hz), and Higher Beta (20-30 Hz). The absolute amplitudes that are plotted in each range represent the average within that range across the entire 6-second delay period. 1) The violin plot shows the distribution of amplitude across each condition; the cluster of similar amplitude values is reflected at the wider (bottom) portion of the violin and the thinner (top) portion with fewer amplitude values. The dashed lines reflect the quartiles of the overall distribution similar to the box plot. 2) The boxplot also shows the distribution of performance for each condition (i.e. the solid lines inside the box align with the dashed lines of the violin plot, representing the quartiles). In addition, the boxplot contains whiskers extend to the minimum and maximum score (1.5 times the median) for each condition and the diamonds reflect outliers (great than 1.5 times the median). Outliers were included in all analyses. 3) In between the violin plot and boxplot is a point plot, which reflects the mean of absolute amplitude, with error bars reflecting the 95% confidence intervals; the confidence intervals were generated with bootstrapping using 1,000 iterations. 4) Finally, the individual dots on the inner-most portion represent a single participants absolute amplitude for a condition (Experiment 1 *n = 24* and Experiment 2 *n* = 20). The gray line connects the dots that represents absolute amplitude value for the corresponding condition (i.e. suppression (left) and rehearsal (right)).

**Supplementary Table 1. Clusters of significantly different absolute amplitude bins between the rehearsal and suppression conditions in Experiment 1.** One hundred and six clusters are listed. Each cluster has a start and stop time during the delay period (between 0-6000 msec), a start and stop frequency (between 4-30 Hz), and lists the electrodes that were involved in the cluster. A positive cluster value is when rehearsal was greater than suppression, and a negative cluster value is the reverse relationship. Electrode names are abbreviated as such: F=frontal, T=temporal, P=parietal, O=occipital, A=anterior, C=central, z= zero; even numbers = right hemisphere, odd numbers = left hemisphere; All channels re-referenced to the average of all channels (Channel name includes “avg”).

**Supplementary Table 2.** **Clusters of significantly different absolute amplitude bins between the rehearsal and suppression conditions in Experiment 2.** Fifteen total clusters are listed. Each cluster has a start and stop time during the delay period (between 0-6000 msec), a start and stop frequency (between 4-30 Hz) and lists the electrodes that were involved in the cluster. A positive cluster value is when rehearsal was greater than suppression, and a negative cluster value is the reverse relationship. Electrode names are abbreviated as such: F=frontal, T=temporal, P=parietal, O=occipital, A=anterior, C=central, z= zero; even numbers = right hemisphere, odd numbers = left hemisphere; All channels re-referenced to the average of all channels (Channel name includes “avg”).

**Supplementary Table 3.** **Clusters of significantly different temporal spectral amplitude bins between the rehearsal and suppression conditions in Experiment 2.** Three clusters which include the same group of neighboring electrodes are listed. Each cluster has a start and stop time during the delay period (between 0-6000 msec), a start and stop frequency (between 4-30 Hz) and lists the electrodes that were involved in the cluster. A positive cluster value is when rehearsal was greater than suppression, and a negative cluster value is the reverse relationship. Electrode names are abbreviated as such: F=frontal, T=temporal, P=parietal, O=occipital, A=anterior, C=central, z= zero; even numbers = right hemisphere, odd numbers = left hemisphere; All channels re-referenced to the average of all channels (Channel name includes “avg”).
