## Supplementary figures and images for "Does rehearsal matter? Left anterior temporal alpha and theta band changes correlate with the beneficial effects of rehearsal on working memory"

### Supplemental Figure 1

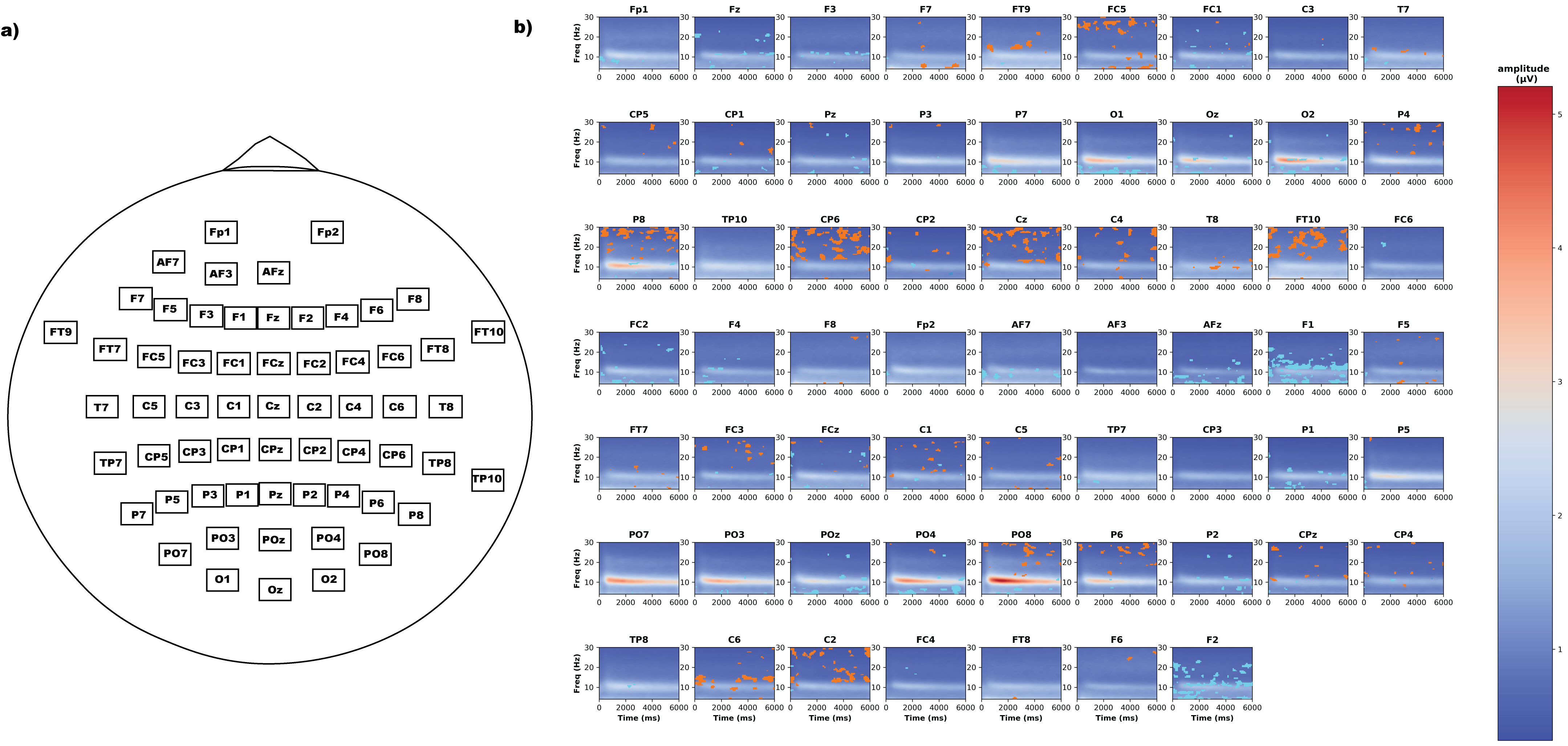

### Supplemental Figure 2

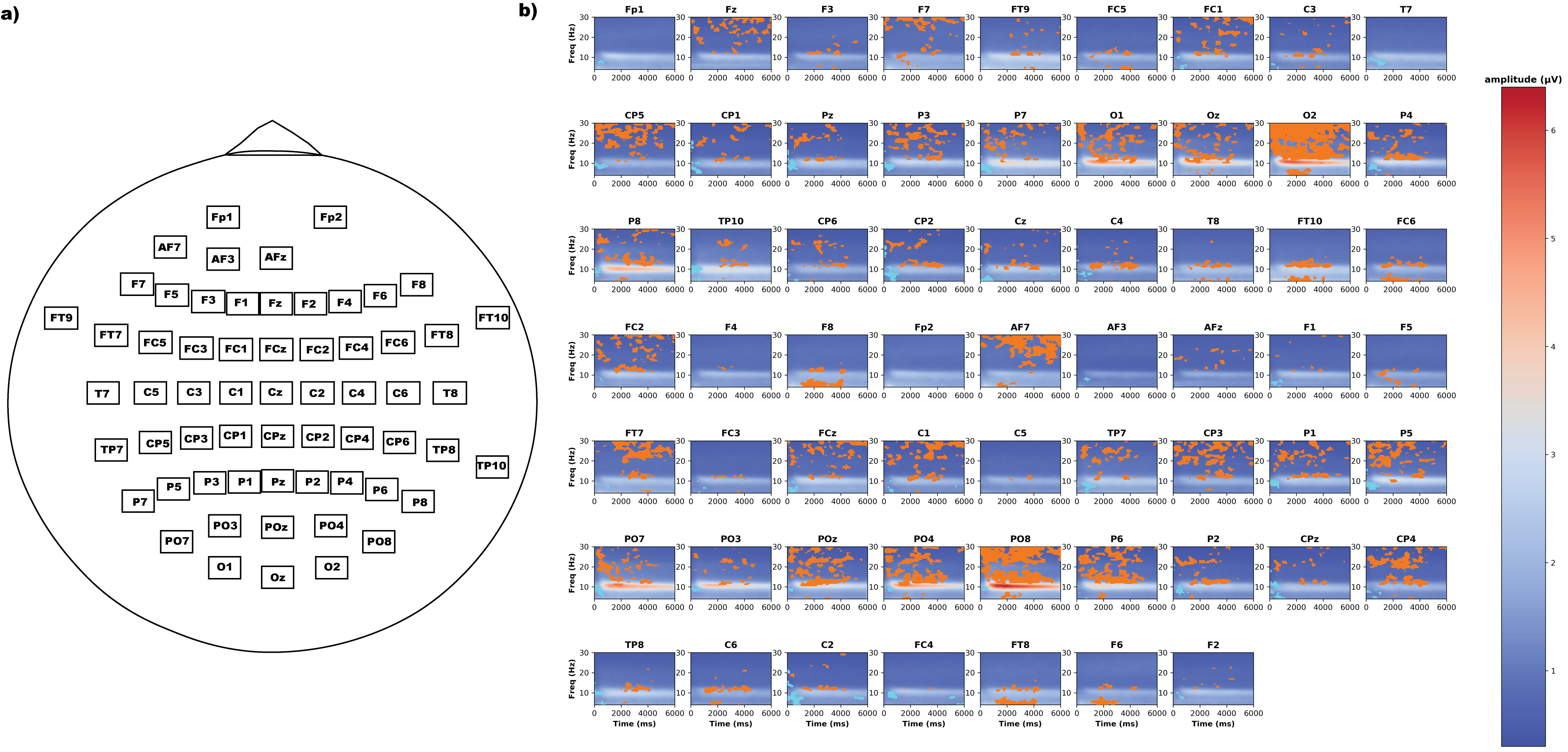

### Supplemental Figure 3

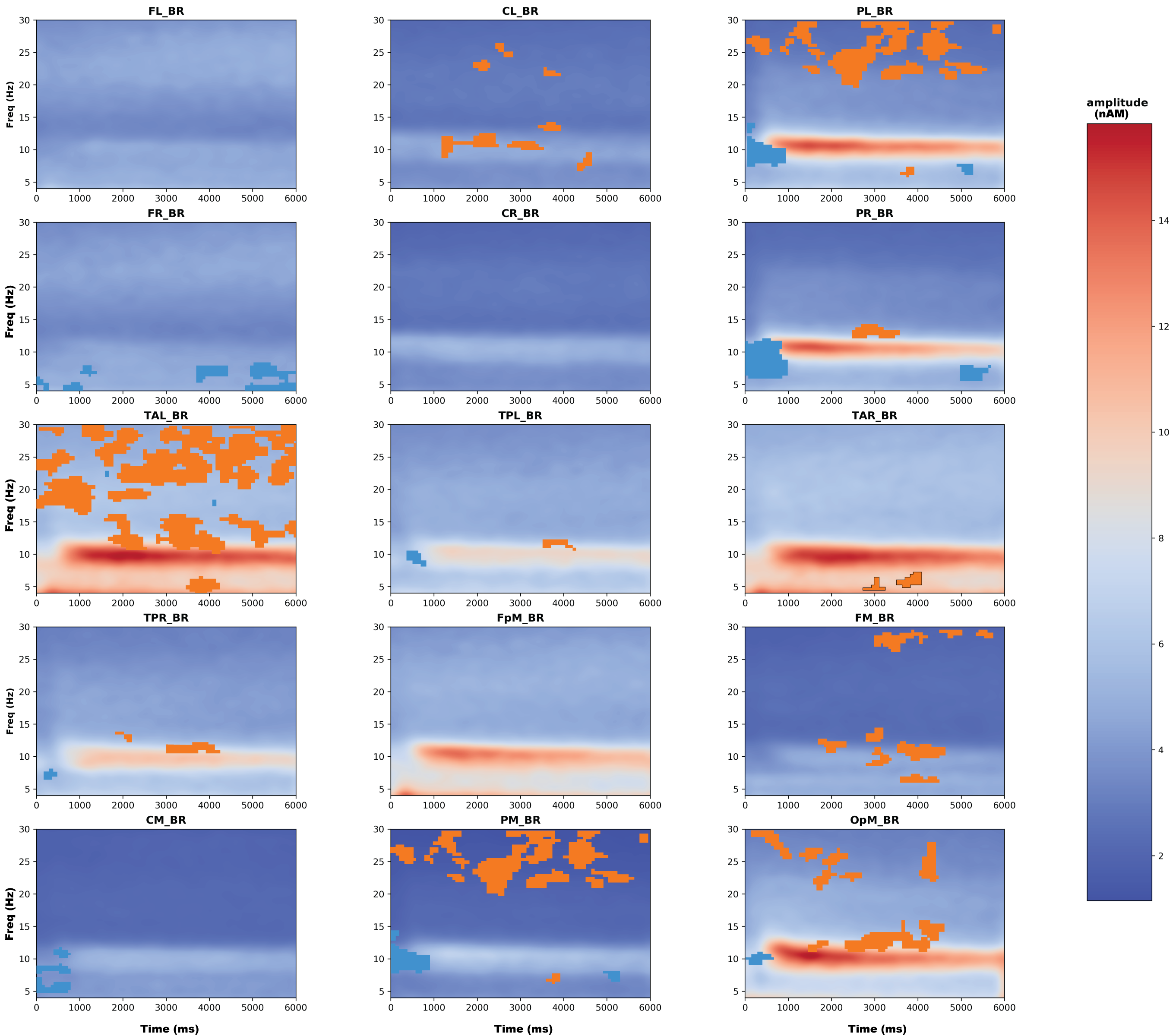

### Supplemental Figure 4

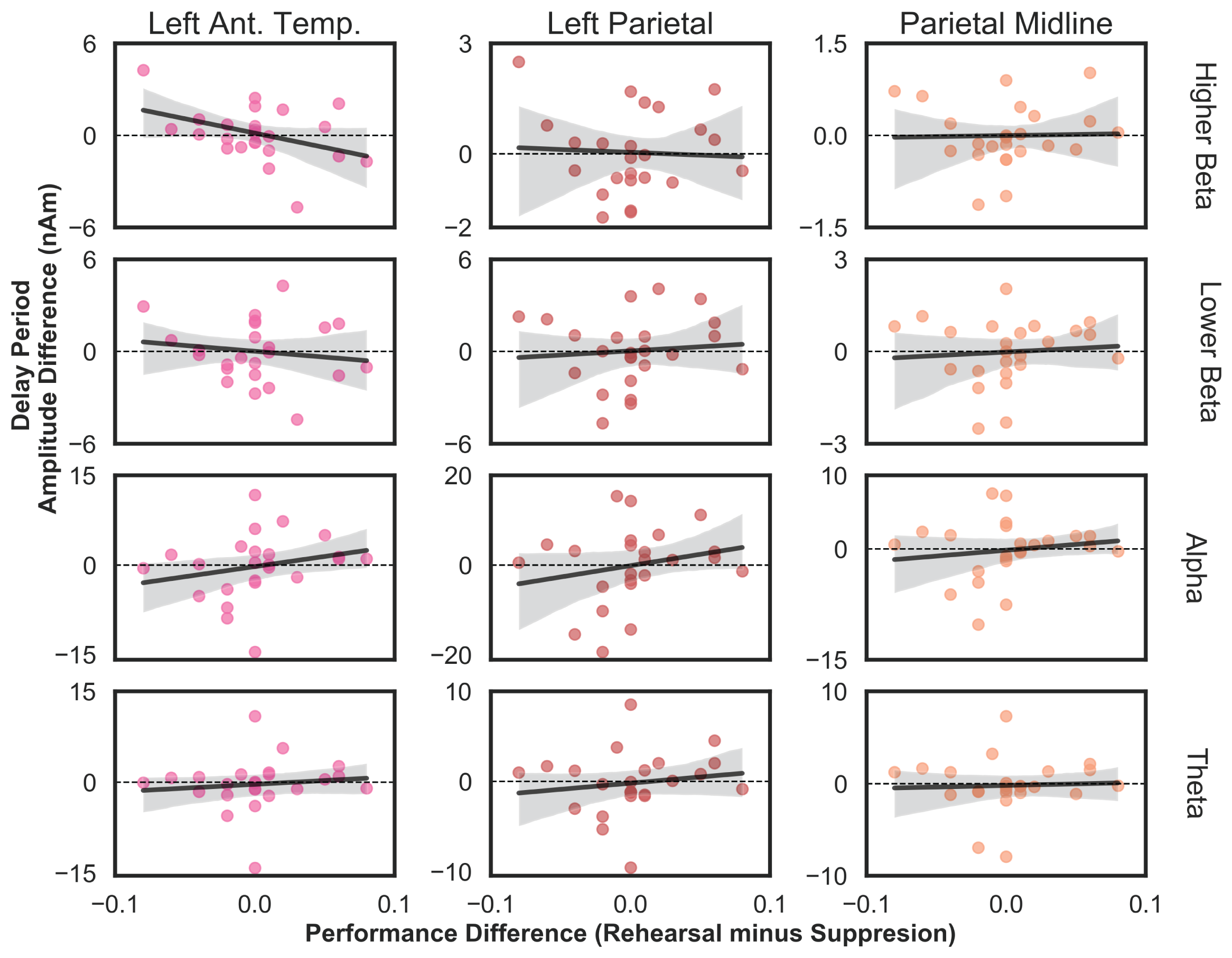

### Supplemental Figure 5

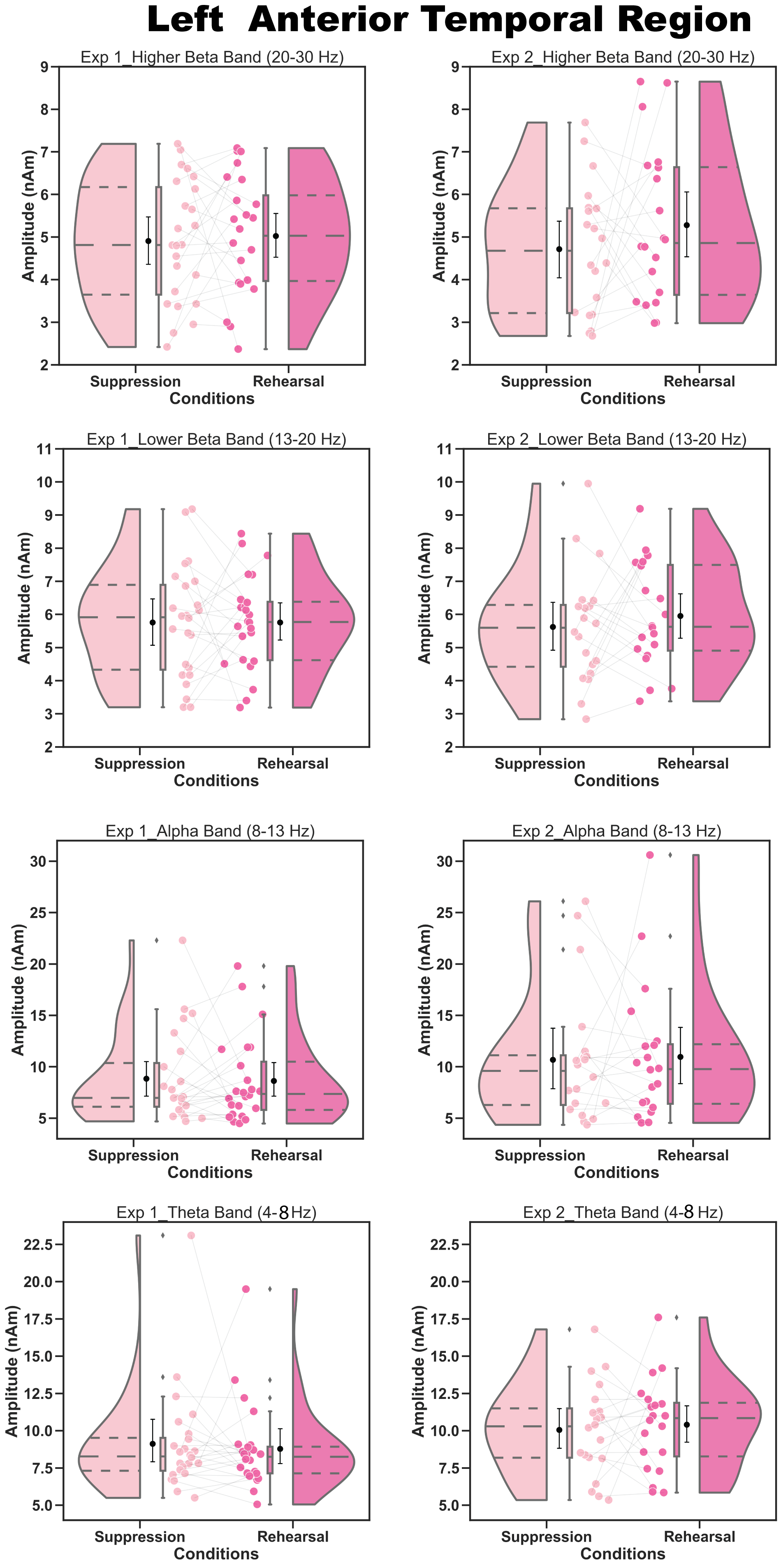

### Supplemental Figure 6

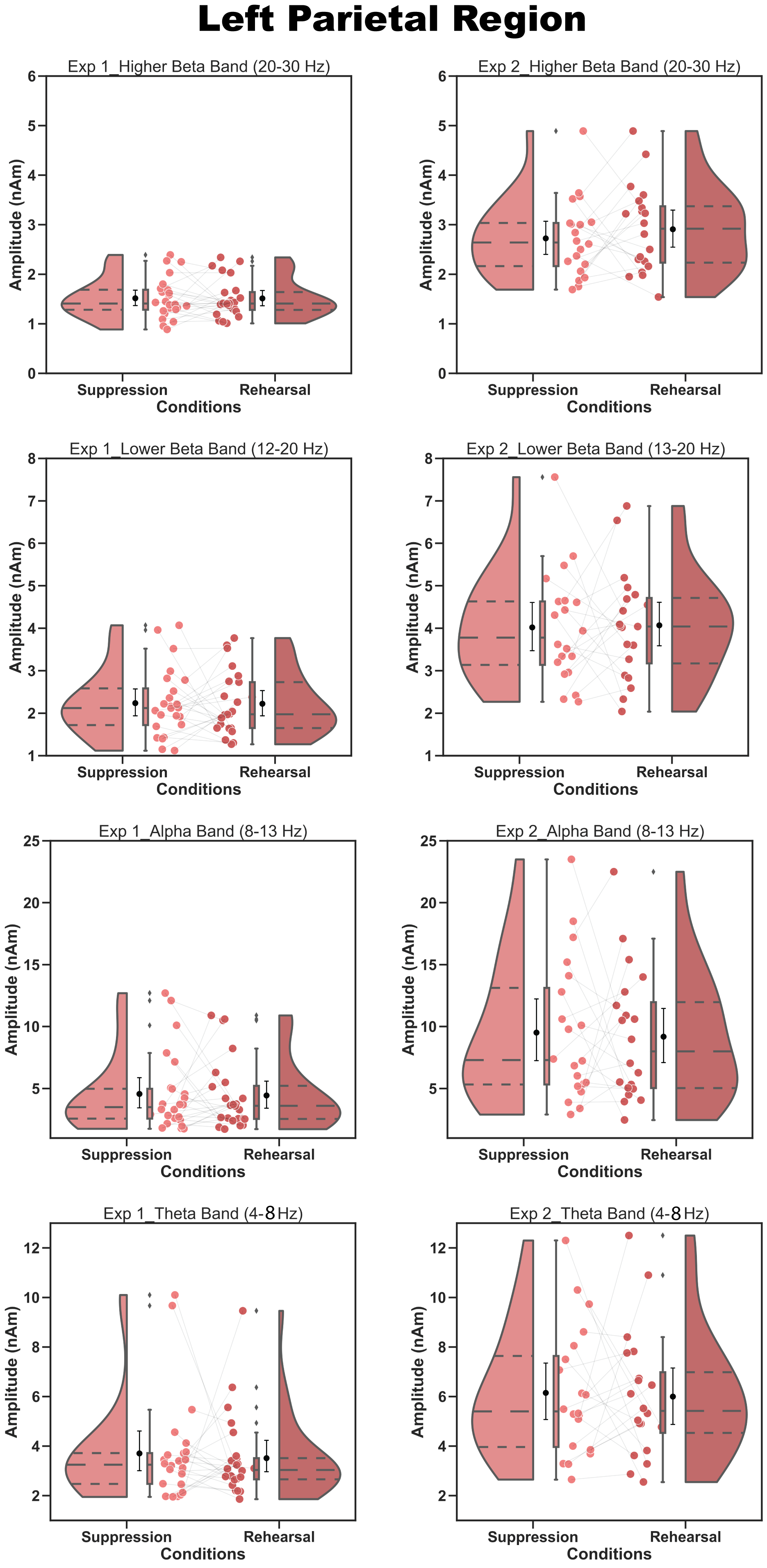

### Supplemental Figure 7

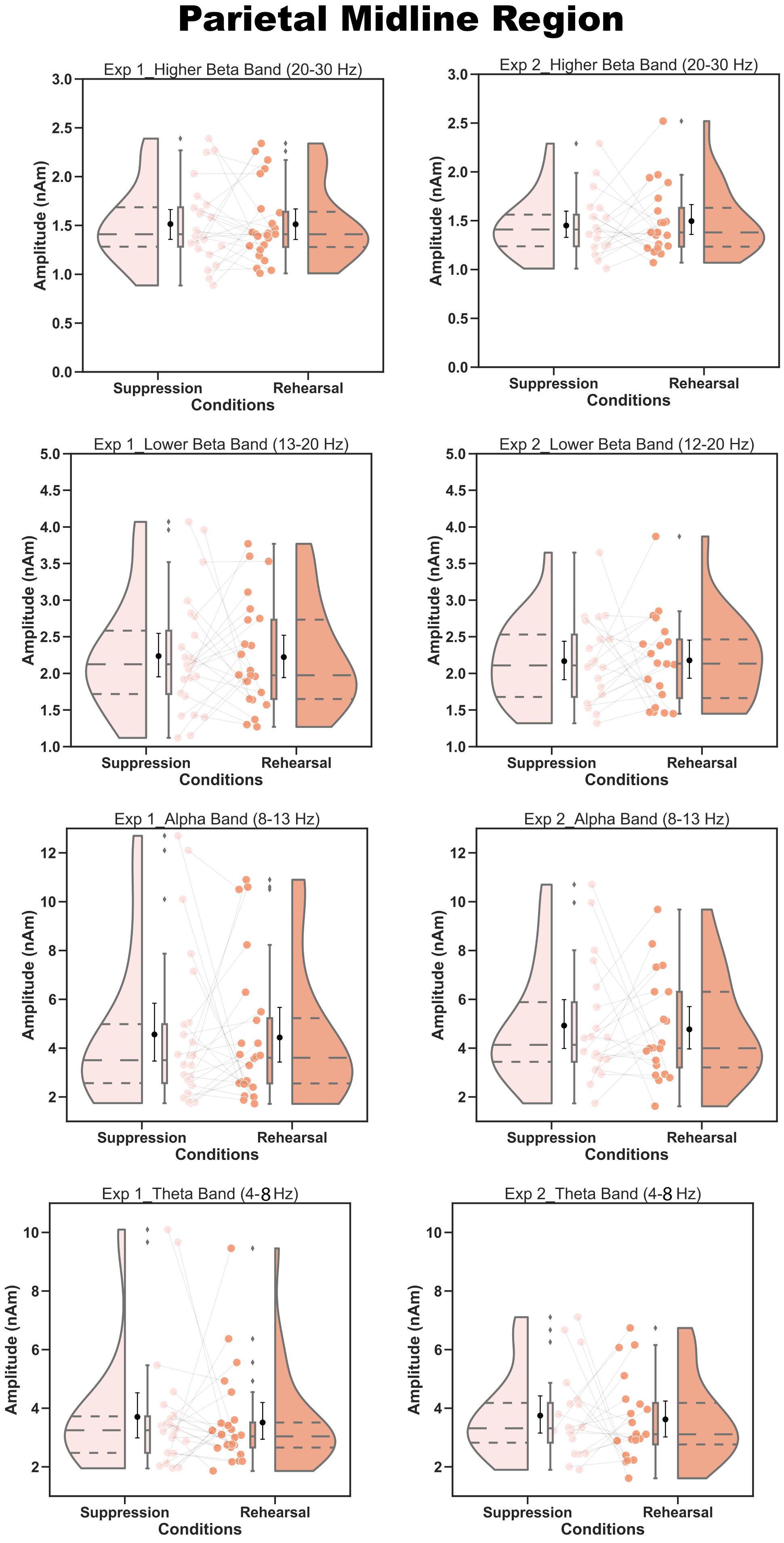
